# A biophysical framework for double-drugging kinases

**DOI:** 10.1101/2023.03.17.533217

**Authors:** C. Kim, H. Ludewig, A. Hadzipasic, S. Kutter, V. Nguyen, D. Kern

## Abstract

Orthosteric inhibition of kinases has been challenging due to the conserved active site architecture of kinases and emergence of resistance mutants. Simultaneous inhibition of distant orthosteric and allosteric sites, which we refer to as “double-drugging”, has recently been shown to be effective in overcoming drug resistance. However, detailed biophysical characterization of the cooperative nature between orthosteric and allosteric modulators has not been undertaken. Here, we provide a quantitative framework for double-drugging of kinases employing isothermal titration calorimetry, Förster resonance energy transfer, coupled-enzyme assays, and X-ray crystallography. We discern positive and negative cooperativity for Aurora A kinase (AurA) and Abelson kinase (Abl) with different combinations of orthosteric and allosteric modulators. We find that a conformational equilibrium shift is the main principle governing this cooperative effect. Notably, for both kinases, we find a synergistic decrease of the required orthosteric and allosteric drug dosages when used in combination to inhibit kinase activities to clinically relevant inhibition levels. X-ray crystal structures of the double-drugged kinase complexes reveal the molecular principles underlying the cooperative nature of double-drugging AurA and Abl with orthosteric and allosteric inhibitors. Finally, we observe the first fully-closed conformation of Abl when bound to a pair of positively cooperative orthosteric and allosteric modulators, shedding light onto the puzzling abnormality of previously solved closed Abl structures. Collectively, our data provide mechanistic and structural insights into rational design and evaluation of double-drugging strategies.

## Introduction

Protein allostery is one of the fundamental regulatory mechanisms involved in various biological processes (1). Specifically, the allosteric regulation of protein kinases has been found essential for signaling cascades. Thus, dysregulation and overexpression of protein kinases are often related to many human diseases, including various cancers. However, due to the highly conserved catalytic site architecture of kinases, specific orthosteric inhibition is often unsuccessful, causing off-target effects (2). In addition, cancers often develop resistant mutations circumventing treatments with orthosteric drugs (3, 4). To overcome these problems, the field has been exploring allosteric sites of kinases for specific and efficacious inhibition (5, 6).

A recently approved allosteric inhibitor of Abelson kinase (Abl), asciminib, has been highly effective in inhibiting Abl *in vitro* and *in vivo* (7–12). Remarkably, dual inhibition of Abl with this allosteric inhibitor combined with the orthosteric inhibitors (including imatinib, nilotinib and ponatinib), which we refer to as “double-drugging”, has been impressively successful in abolishing the emergence of resistant mutants for Abl (12–14). Considering this clinical benefit, this approach has been applied to inhibit other targets such as EGFR kinase and SHP2 phosphatase (15, 16). However, the biophysical mechanisms underlying double-drugging of distant orthosteric and allosteric sites have not been well studied.

Herein, we provide the quantitative framework for double-drugging using two targets: Aurora A kinase (AurA) and Abelson kinase (Abl). Both kinases participate in various cellular pathways, and their dysregulation results in a multitude of cancers, such as breast cancer and leukemia (17–19). Common obstacles faced by orthosteric inhibitors for AurA and Abl include cytotoxicity, off-target effects, and emergence of resistance mutants (3, 4, 20, 21). For both systems, we exploit a rational selection of ligands to probe positive and negative cooperativity between remote orthosteric and allosteric sites using isothermal titration calorimetry (ITC), Förster resonance energy transfer (FRET) and coupled-enzyme assays. We find that both orthosteric and allosteric ligands exhibit preferred binding to the active or inactive states, and that cooperativity occurs by shifting this active-inactive conformational equilibrium through long-range allosteric networks that are encoded for natural regulation of those kinases. X-ray crystal structures of the double-drugged complexes shed light onto the atomistic mechanisms of cooperativity. After we determine negative cooperativity for the double-drug combination used by Novartis in their clinical trials, we rationally chose a different orthosteric inhibitor, Src inhibitor 1 (SKI), for positive cooperativity with asciminib. This new double-drug combination forms a unique ternary complex, revealing the first fully-closed Abl structure.

## Results

### Cooperative binding between orthosteric and allosteric modulators of AurA

In solution, AurA exists in a conformational equilibrium between active and inactive states (22–24). We previously designed monobodies (Mbs) that are fully selective allosteric modulators, which bind to the natural allosteric regulatory site of AurA on the N-terminal lobe (N-lobe), the binding site for the natural coactivator protein TPX2 (25, 26). Different monobodies either act as activators or inhibitors depending on how they shift the active/inactive conformational equilibrium of AurA (25). To achieve double-drugging on AurA, we combined these Mbs with the orthosteric inhibitor danusertib (PHA739358) that tightly binds to AurA (IC_50_ = 13 nM, *K*_i_ = 0.87 ± 1.44 nM (27)). Since it has been shown that danusertib preferentially binds to the inactive conformation of AurA (22, 28), we hypothesized that inhibiting Mbs would bind tighter to AurA when in complex with the orthosteric inhibitor danusertib. Conversely, binding of activating Mbs to AurA should be weakened in the presence of danusertib **(Fig. 1*A*)**.

**Figure 1:**
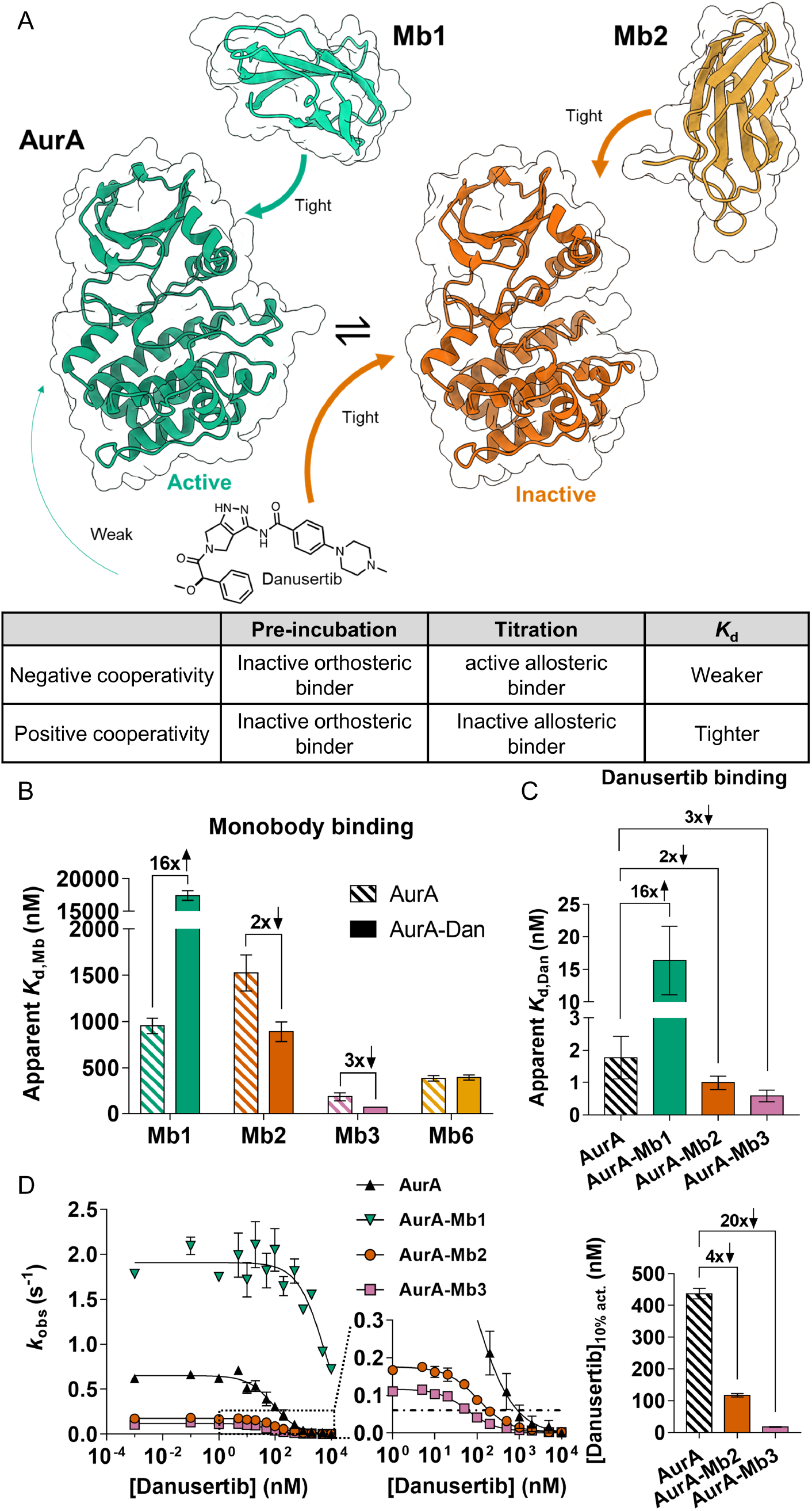
Double-drugging of AurA kinase with orthosteric drug danusertib and different allosteric modulators. (A) Schematic representation of active/inactive equilibrium of AurA (green, PDB-ID: 5G15 and orange, PDB-ID: 6C83 (25)). Arrows indicate binding of danusertib and Mbs to their preferred AurA conformations. The table represents the rationale of positive and negative cooperativity for double-drugging of AurA. (B) Effect of preincubation with danusertib on observed dissociation constants (apparent *K*_d_) of different monobodies measured by ITC. Activating monobody Mb1 shows 16-fold negative cooperativity, while inhibiting monobodies, Mb2 and Mb3 show 2-fold and 3-fold positive cooperativity, respectively. Mb6 binding is not affected by the presence of danusertib **(Fig. S3)**. (C) Reversal of pre-incubation order during affinity measurements shows identical cooperativities for orthosteric/allosteric ligand combinations **(Fig. S1)**. Errors in (B,C) represent 68.3% confidence interval of the fit of the data (± 1 s.d.). (D) Kinase inhibition curves of AurA and AurA in the presence of saturating concentrations of Mb1, Mb2 and Mb3 as a function of danusertib concentration. Enzyme assays were conducted (*n* = 2, mean ± s.d.m.) under *k*_cat_/*K*_m_ condition with 3 mM Lats2 peptide, measuring observed activity (*k*_obs_). With inhibiting Mb2 and Mb3, 4-fold and 20-fold lower concentrations of danusertib, respectively, are required to inhibit to 10% residual AurA activity ([Danusertib]_10% act_., and dashed line). Errors in this bar graph were determined by jackknifing the inhibition curve data.

Aligning with our hypothesis, we find that the binding affinity of activating monobody (Mb1) to AurA weakens 16-fold when AurA is pre-saturated with danusertib **(Fig. 1*B*)**. To test whether the binding of Mb1 and danusertib are mutually exclusive, we repeated this experiment by pre-incubating AurA with a higher concentration of danusertib **(Fig. S1*E*)**. Identical Mb1 binding affinities, independent of saturating danusertib concentrations, reveal that the simultaneous binding of Mb1 and danusertib to AurA is possible. Thus, we reason that this 16-fold negative cooperativity for Mb1 binding arises from a conformational equilibrium shift of AurA to the inactive state induced by danusertib.

To achieve desired positive cooperativity between allosteric and orthosteric binders to AurA, we chose the inhibiting monobodies Mb2 and Mb3, because i) Mb2 is an inhibiting monobody that we had obtained an X-ray crystal structure in complex with AurA, ii) Mb3 exhibits larger inhibition than Mb2, and iii) AurA-Mb3 complex exists in a monomeric form unlike the dimeric AurA-Mb2 complex (25). We indeed measure a 2-fold tighter binding of Mb2 to the AurA-danusertib complex compared to apo AurA **(Fig. 1*B*)**. Using the equilibrium constant for active/inactive states of AurA previously determined (*K*_eq_= 0.67 (22)) and assuming identical affinities of Mb2 to the inactive states of apo-or danusertib-bound AurA, we fit our apparent affinities to a reversible two-state allosteric model. We find that the 2-fold positive cooperativity can be explained solely by the shift in the conformational equilibrium **(Fig. S2)**. Thus, a further increase in positive cooperativity would only be possible if the Mb affinity was tighter to the inactive state of the AurA-danusertib complex than to the inactive state of apo AurA. We indeed observed a 3-fold positive cooperativity for Mb3 with danusertib **(Fig. 1*B*)**. We speculate that this increased affinity of Mb3 to the AurA-danusertib complex compared to apo AurA could result from favorable interactions with a closed activation loop, since danusertib binding shifts the equilibrium of the activation loop towards such conformation (23, 28).

To confirm whether the mechanism of cooperativity between Mbs and danusertib follows a classic allosteric model, we tested binding of Mb6 to the AurA-danusertib complex. Despite high affinity, Mb6 binding does not change AurA’s activity, implying that Mb6 binding does not shift the active/inactive conformational equilibrium of AurA (25). Indeed, the binding affinity of Mb6 to AurA is not changed in the presence of danusertib **(Fig. 1*B*, Fig. S3)**.

For a reversible two-state allosteric model, the same fold-change of cooperativity must be observed when reversing the order of binding. To measure changes in the affinity of danusertib upon Mb binding, we had to employ competitive replacement ITC with adenosine 5’-(α, β-methylene) diphosphate (AMPCP), since danusertib binds too tightly to AurA for direct measurement. Indeed, the measured cooperativities are matching quantitatively regardless of the binding order **(Fig. 1*C*)**.

### Double-drugging of AurA lowers inhibitor concentration needed for efficacious inhibition

Next, we probed biological relevance of these observed cooperativities by measuring the inhibition of AurA kinase activity using Lats2 peptide as a substrate with cellular ATP concentrations. Pre-incubation of inhibiting Mbs resulted in a vastly decreased amount of danusertib required to cause 90% inhibition of AurA activity **(Fig. 1*D*)**. This combined inhibition effect is the direct consequence of positive cooperativity. For instance, Mb3, which displays a larger degree of positive cooperativity than Mb2, causes a larger reduction in required amount of danusertib for effective inhibition.

### X-ray crystal structures of ternary complexes: AurA-danusertib-Mb1 and AurA-danusertib-Mb2

We solved X-ray crystal structures of double-drugged AurA complexes to further understand the structural features responsible for the positive and negative cooperativity between Mbs and danusertib **(Table S1)**. The complex of AurA-danusertib-Mb1 displays hallmarks of an active kinase, such as an intact regulatory spine, the α-C helix in the “in” position and the “DFG-in” state **(Fig. 2*A*)**. However, we note that in contrast to the AurA-AMPPCP-Mb1 structure (PDB-ID: 5G15), D274 is rotated away from danusertib to avoid a steric clash with the terminal phenyl ring of danusertib **(Fig. S4*A*)**. This crystal structure corroborates the capability of danusertib to bind to the active conformation of AurA as we had tested biochemically **(Fig. S1*B*)**.

**Figure 2:**
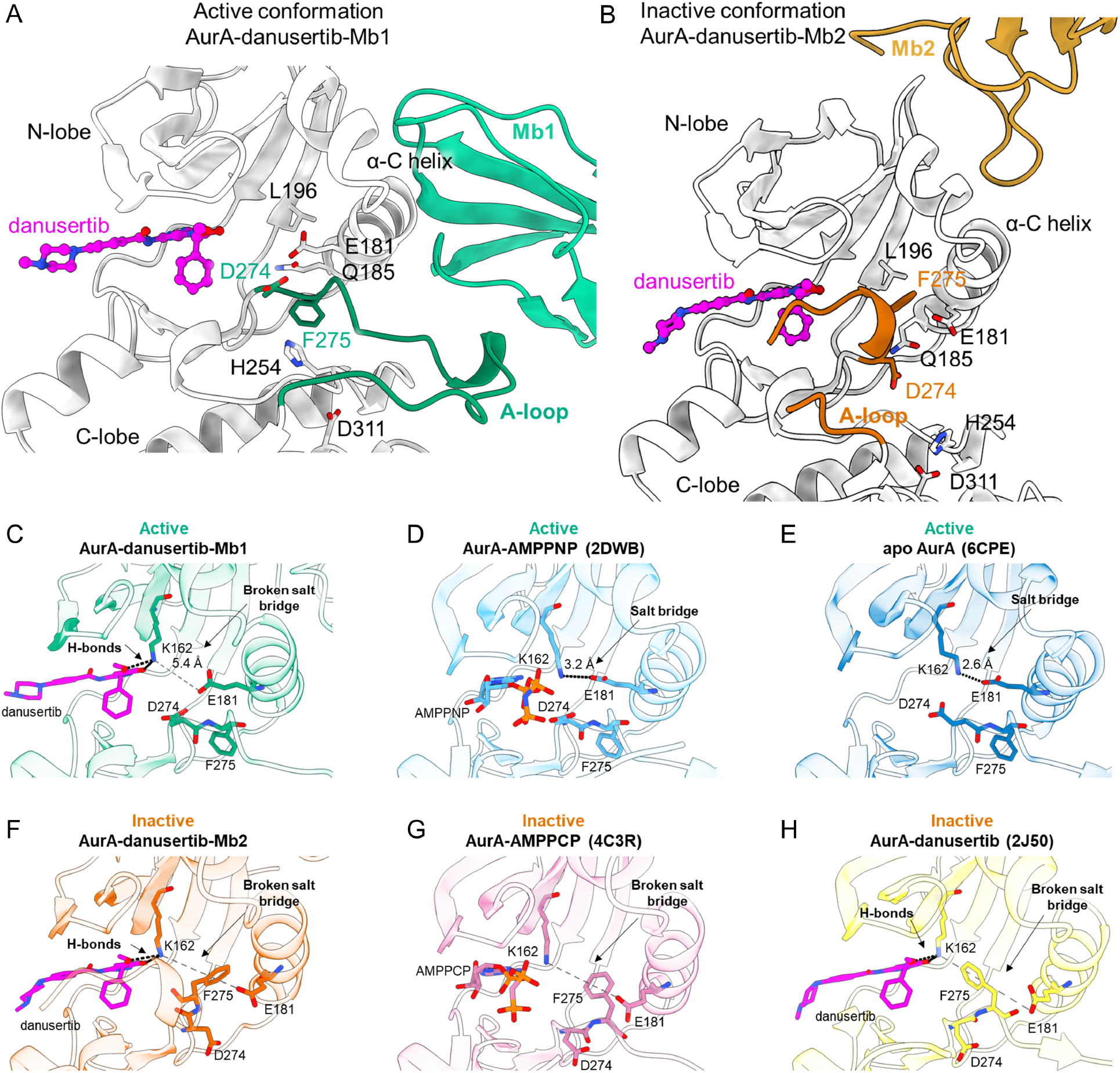
Proposed molecular mechanism for negative and positive cooperativity for double drugging AurA with danusertib in combination with Mb1 and Mb2, respectively. (A-B) Zoom-in of X-ray crystal structures of AurA (grey) complexed with danusertib and either Mb1 (A, green) or Mb2 (B, gold). An intact regulatory spine, DFG-in conformation and extended activation loop in (A) is contrasted to a broken regulatory spine, DFG-out, and closed activation loop in (B). This closed activation loop provides additional hydrophobic interaction to the terminal ring of danusertib. (C-H) Orthosteric binding sites for six different AurA states reveal why danusertib has higher affinity for inactive AurA (22, 27, 31). K162 and E181, which form the canonical salt-bridge in active AurA, and D274 and F275 (DFG-motif), are shown in stick representation. (D-E) While K162-E181 salt-bridge is established in an active AurA conformations, (C) this salt-bridge is broken in AurA-danusertib-Mb1 structure as K162 interacts with danusertib. (F-H) In the inactive AurA conformations, DFG-out F275 is positioned between K162 and E181, physically blocking the salt-bridge interaction, thereby pre-positioning K162 for danusertib binding. (A-H) Oxygen, nitrogen and phosphorous atoms are colored in red, blue and orange, respectively. Carbon atoms are colored according to their respective protein cartoon.

The most interesting structure for “double-drugging” with maximal inhibition is of course the ternary complex of AurA-danusertib-Mb2. Like AurA-AMPPCP-Mb2 (PDB-ID: 6C83), the new ternary complex displays features of an inactive kinase: α-C helix “out”, “DFG-out” as well as both a broken regulatory spine and a broken canonical salt-bridge (K162-E181) **(Fig. 2*B*)**. This is expected due to the conformational equilibrium shift caused by Mb2 binding and the preferential binding of danusertib to inactive AurA. Furthermore, the activation loop is fully shifted towards the active site, providing additional hydrophobic interactions to the terminal ring of danusertib **(Fig. 2*B*)**. This shifted activation loop is a major structural feature of an inactive AurA (23, 28), as observed in AurA bound to the orthosteric inhibitor MLN8054 (PDB-ID: 2WTV) (29). The structure of AurA-AMPPCP-Mb2 (PDB-ID: 6C83) displays a similar activation loop, however, with an extended portion being disordered (residues 276-290) to circumvent clashing with the β- and γ-phosphate groups of AMPPCP **(Fig. S4*B*)**. The binary complex between AurA and danusertib (PDB-ID: 2J50) did not exhibit such a shift in the activation loop. However, it is unclear whether the activation loop conformation in the AurA-danusertib structure reflects the solution state since the activation loop is directly involved in crystal contacts **(Fig. S5)**.

Our crystal structures and ITC experiments showed that danusertib can bind to both the AurA-Mb2 and AurA-Mb1 complexes. To reveal why danusertib, however, binds with much higher affinity to AurA-Mb2 than AurA-Mb1 (there is no steric hindrance), we scrutinized the thermodynamic parameters of our ITC studies on danusertib binding to different AurA-Mb complexes **(Fig. S1*A-D*)**. We find that the enthalpy for danusertib binding to AurA-Mb1 is reduced by 22.8 kJ/mol compared to AurA-Mb2, which approximates the equivalence of one salt-bridge (12.6-20.9 kJ/mol (30)). The canonical salt-bridge between K162 and E181 is a feature of an active AurA, in both its apo form and bound to AMPPNP (PDB: 6CPE and 2DWB, respectively) (22) **(Fig. 2*DE***). In contrast, the ternary complex of AurA-danusertib-Mb1 displays a broken salt-bridge, as K162 interacts now with danusertib, while maintaining the α-C helix in the “in” position **(Fig. 2*C*)**. Thus, we propose that the K162-E181 salt-bridge in AurA-Mb1 must be broken for danusertib binding, as reflected by the lowered binding enthalpy. To confirm that apo AurA-Mb1 complex establishes the K162-E181 salt-bridge, we deleted danusertib from the structure of AurA-Mb1-danusertib and carried out molecular dynamics simulations in triplicate. We observed that K162-E181 indeed forms this saltbridge on average 80.8% in a 10 ns simulation **(Fig. S6*A*)**. However, in the presence of danusertib, we observe K162 to rather form a hydrogen bond with *O*-27 of danusertib’s methoxy moiety than with E181 in the MD simulation, which is the state sampled in our crystal structure as well **(Fig. 2*C*, Fig. S6*BC*)**.

In the inactive conformations of AurA, the broken K162-E181 salt-bridge stems from the α-C helix and DFG-motif being positioned in the “out” conformation, such that F275 positions between K162 and E181 (PDB: 4C3R and 2J50) (27, 31) **(Fig. 2*GH*).** Thus, we propose that K162 in inactive AurA conformations, such as AurA-Mb2 complex, is pre-positioned for danusertib binding **(Fig. 2*F*)**, which results in the tighter binding of danusertib to the inactive state of AurA.

### Cooperative effect of imatinib and asciminib binding on Abl

Intrigued by our mechanistic insights into double-drugging of AurA, we turned to Abl, the only target currently in clinical trials for double-drugging. It has been shown that the combination of the orthosteric inhibitor imatinib and the allosteric inhibitor asciminib abolishes the emergence of resistance mutations (12, 32–34), an impressive breakthrough. Therefore, Abl embodies a powerful target to delineate the biophysical constraints, or “framework”, for successful double-drugging. Since the quantitative biophysical parameters for this drug combination are not known, we set out to biophysically investigate the cooperativity and modulation of Abl’s open/closed conformational equilibrium first using this exact combination of orthosteric and allosteric inhibitors.

Abl exists in a conformational equilibrium between open, active and closed, inactive conformations (32, 35, 36) **(Fig. 3*A*)**. In the open conformation, the regulatory domains are elongated so that the SH2 domain moves onto the N-lobe of the kinase domain, forming a “top-hat” conformation (35). In the closed conformation, the regulatory domains tightly interact with the kinase domain, SH3:N-lobe and SH2:C-lobe, the latter facilitated by the bent C-terminal α-I helix (12, 35, 37) **(Fig. 3*A*)**. This conformational equilibrium is susceptible to modulation by single agents such as imatinib and asciminib (12, 32, 38).

**Figure 3:**
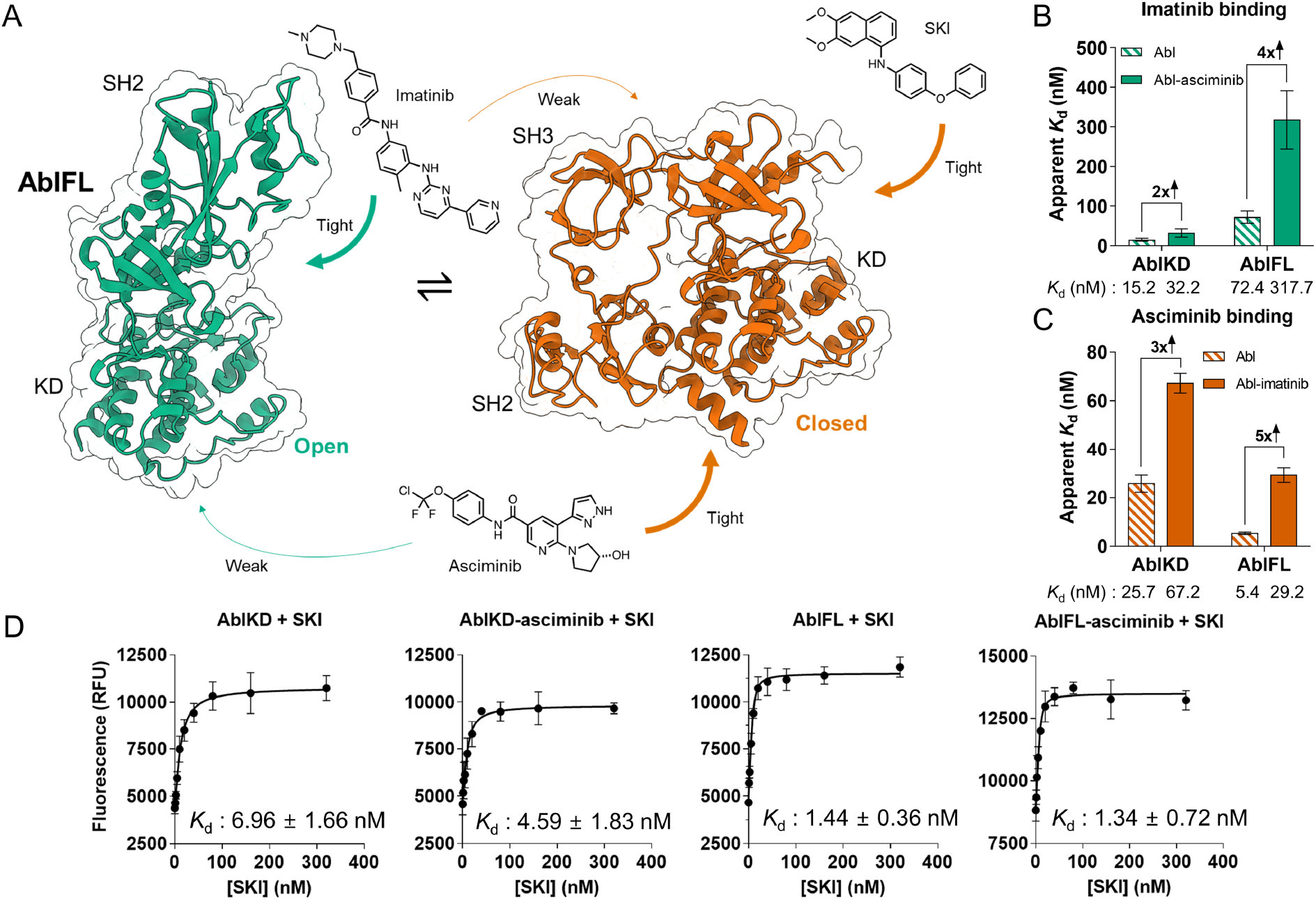
Double-drugging of Abl kinase. (A) Schematic representation of conformational equilibrium in Abl kinase. Arrows indicate binding of orthosteric inhibitors imatinib and SKI, and allosteric inhibitor asciminib to preferred Abl conformations. X-ray crystal structures from PDB-ID: 1OPL (green) and PDB-ID: 5MO4 (red) were used for open and closed Abl structures, respectively (12, 35). Note that the SH3 domain in the open structure is missing due to lacking electron density. (B) With ITC experiments, we observe 2- and 4-fold negative cooperativity for open-conformation binder imatinib when AblKD and AblFL, respectively, are pre-incubated with closed-conformation binder asciminib **(Fig. S8)**. (C) Matching fold-change of negative cooperativity is observed reversing the order of modulators used in pre-incubation and titration, using ITC **(Fig. S8)**. (B-C) Errors in the bar graphs represent 68.3% confidence interval of the fit of the data (± 1 s.d.). (D) FRET experiments to detect SKI binding (10 nM of enzyme in all experiments. Data (*n* = 3, mean ± s.d.m.) have been fitted to quadratic binding equation. Unlike imatinib, SKI binds tighter to AblKD-asciminib and AblFL than to AblKD. In AblKD, asciminib exhibits small positive cooperativity with the binding of SKI. *K*_d_ errors are standard error of the fit.

In ITC experiments, we find that imatinib binds 5-fold tighter to AblKD, which exists exclusively in the open conformation, than to AblFL **(Fig. 3*B*)**. This confirms imatinib’s preferential binding to the open state of Abl (32). In full agreement with this model, imatinib binds to AblFL with a 4-fold decreased affinity in the presence of asciminib, since asciminib shifts the equilibrium to the closed state (12) (**Fig. 3*B*)**. We conclude that this 4-fold negative cooperativity between imatinib and asciminib stems from a shift in the conformational equilibrium of Abl, where both drugs preferentially bind to the open and closed conformation, respectively. Akin to AurA, pre-incubation of AblFL with increased concentration of asciminib did not result in a weakened imatinib affinity, confirming the simultaneous binding of the two inhibitors **(Fig. S8*B*)**. Surprisingly, we found that imatinib and asciminib display a 2-fold negative cooperativity for AblKD **(Fig 3*B*)**. This implies the presence of an additional conformational equilibrium within the kinase domain itself (39), and that asciminib and imatinib shift this equilibrium in opposite directions. We refer herein to the asciminib-favoring conformation as the “closing-competent” conformation of AblKD.

Importantly, we measure identical negative cooperativities between imatinib and asciminib on both AblKD and AblFL, regardless of binding order, within the range of errors **(Fig. 3*C*, Fig. S8*E*).** Due to the tight binding of asciminib, its affinity was measured via competitive replacement ITC using N-Myr peptide as a weak-binding ligand (12, 40). Collectively, we conclude that the binding of imatinib to the orthosteric site and asciminib to the allosteric site in AblFL follow a two-state allosteric model, in which the two drugs favor the closed and open conformation, respectively.

### Positive cooperativity between SKI and asciminib on Abl

Considering the negative cooperativity between imatinib and asciminib described by our ITC experiments, we wanted to rationally select an orthosteric inhibitor that exhibits positive cooperativity with asciminib. We chose Src inhibitor 1 (SKI), an orthosteric inhibitor that tightly binds to Src kinase (IC_50_ = 44 nM) (41,42), because Bannister *et al*. recently measured that SKI preferentially binds to the α-C helix out, inactive conformation of Src kinase (Unpublished data, Bannister *et al.*). We hypothesized SKI would have the same effect on Abl due to its structural homology with Src kinase. Since SKI binding to Abl did not result in a detectable heat change in ITC, we turned to FRET experiments to quantify this interaction **(Fig. 3*D*, Fig. S9)**. SKI indeed binds preferentially to the closed conformation of AblFL, as seen by the 5-fold tighter binding of SKI to AblFL than to AblKD. Furthermore, we observe a modest positive cooperativity between SKI and asciminib binding in AblKD, indicating that SKI binds to the “closing-competent” conformation induced by asciminib. Unexpectedly, we did not find a difference between the binding affinities of SKI to AblFL and AblFL-asciminib. We interpret this result as evidence that the conformational equilibrium of apo AblFL is already far shifted to the closed conformation. Hence, the binding of asciminib had no effect on this equilibrium. This is, in fact, in agreement with a NMR study by Grzesiek and colleagues reporting overlapping chemical shifts between apo and GNF-5 (a predecessor of asciminib) bound AblFL for open/closed equilibrium markers (32).

### Effect of orthosteric and allosteric modulators on Abl activity

Interestingly, it had been reported that allosteric inhibitors of Abl other than asciminib (such as GNF-2, GNF-5, myristate and myristoyl-peptide) actually do not inhibit the catalytic activity despite binding to AblKD (40, 43, 44). However, with ITC, we observed that asciminib shifts the conformation of AblKD to the “closing-competent” conformation **(Fig. 3*BC*)**. Is this “closing-competent” conformation of the kinase domain a catalytically inactive state of Abl? Inhibition curves of AblKD generated using a coupled-enzyme assay with Srctide as substrate and asciminib as an inhibitor reveal 30% inhibition at saturating asciminib concentration **(Fig. 4*A*)**. We conclude that the closing-competent conformation of the kinase domain is indeed catalytically inactive and that asciminib shifts the conformational equilibrium of AblKD to be 30% in this conformation by binding to the C-lobe and allosteric propagation to the orthosteric site. This model also reconciles the moderate synergistic effect of SKI and asciminib binding to AblKD **(Fig. 3*D*)**. We note that this unique allosteric propagation by asciminib could contribute to its increased potency relative to other myristate pocket binders. Most importantly, and stressing the importance of studying full-length kinases in drug development, asciminib causes a 93% inhibition of AblFL at saturating concentration **(Fig. 4*A*)**. This vastly increased inhibition is caused by the closing of the regulatory domains leading to an inactive kinase.

**Figure 4:**
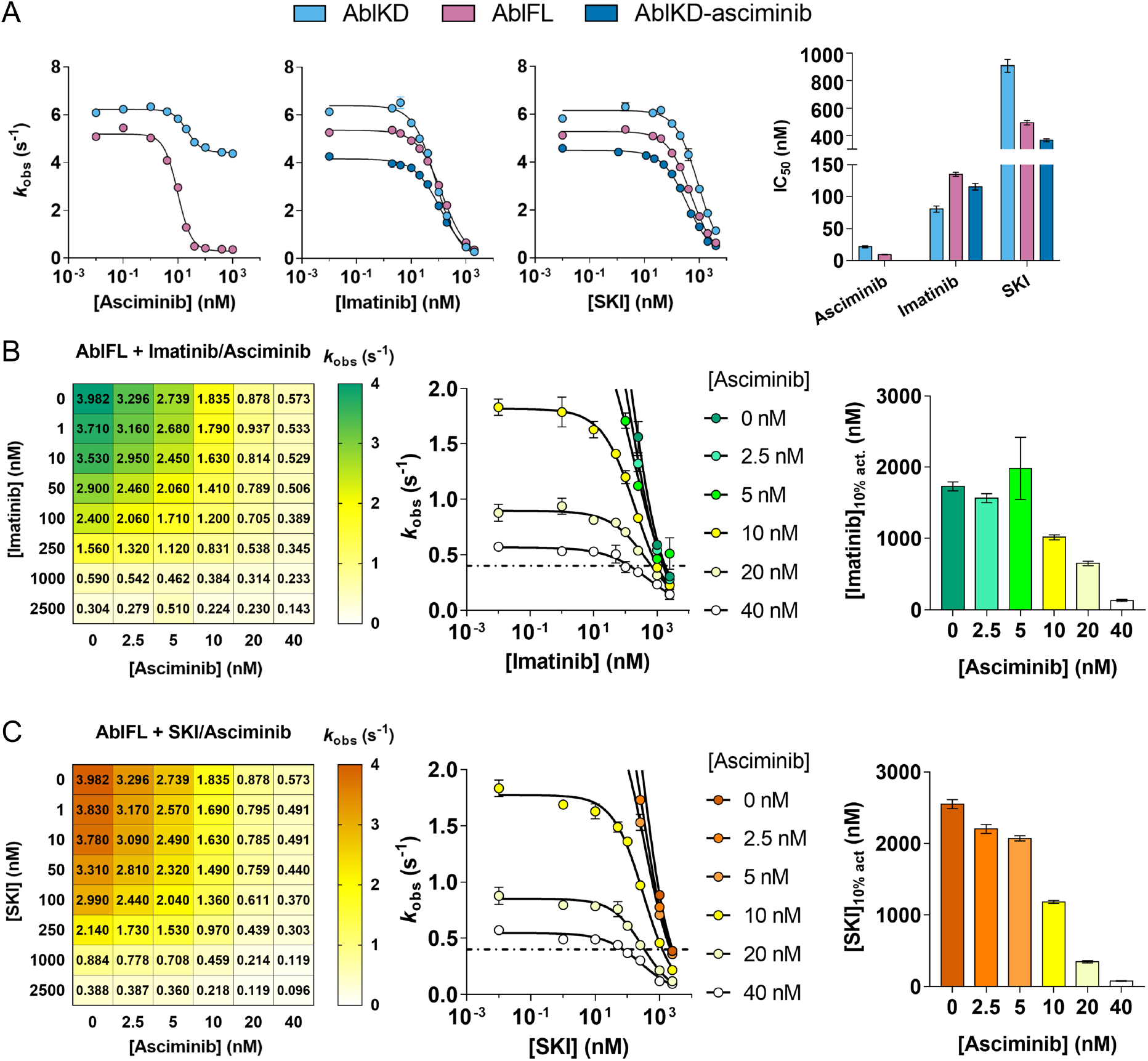
Catalytic activities of Abl kinase under double-drugging conditions. (A) Inhibition curves of AblKD and AblFL with asciminib, imatinib, and SKI. IC_50_ values shift according to the favored binding conformations of the corresponding inhibitor. (B-C) *In vitro* synergy studies using (B) imatinib and asciminib (C) or SKI to examine cooperativity between both inhibitors for AblFL activity. Left: Numbers in the grid represent *k*_obs_ with respective concentrations of inhibitor combinations. Middle and right: graphic representation of the data to illustrate inhibitor concentration needed to achieve 10% residual kinase activity ([Imatinib]_10% act_. and [SKI]_10% act_, dashed line). Note that SKI required for 10% residual kinase activity decreases with increasing asciminib, especially with higher fold-change than that observed for imatinib. All assays were measured (*n* = 4 for 0 nM orthosteric inhibitor, *n* = 2 for all other assay, mean ± s.d.m.) under *k*_cat_/*K*_m_ condition with 2 mM Srctide. Errors in IC_50_ are standard errors of the fit. Errors in the bar graphs were determined by jackknifing the inhibition curve data.

Next, we quantified the inhibition of AblKD and AblFL by the two orthosteric inhibitors imatinib and SKI. In agreement with our affinity measurements **(Fig. 3*B-D*)**, imatinib exhibited a lower IC_50_ for AblKD than for AblFL, while SKI exhibited a higher IC_50_ for AblKD than for AblFL. Second, pre-incubation of AblKD with asciminib increased the IC_50_ for imatinib. This negative cooperativity arises from binding preferences of imatinib and asciminib to opposite conformations. In contrast, double-drugging of AblKD with SKI and asciminib resulted in a reduced IC_50_ since both have a binding preference to the same, “closing competent” conformation, highlighting their positive cooperativity **(Fig. 4*A*)**.

We note that SKI’s IC_50_ is higher than imatinib’s IC_50_ with respect to AblKD, AblFL and AblKD-asciminib **(Fig. 4*A*)**, whereas this trend is reversed in our binding experiments **(Fig. 3*BD*)**. We ascribe this discrepancy to the presence of ATP in the coupled-enzyme assay: AMPPCP binds 2-fold tighter to AblKD than AblFL **(Fig. S11)**. Thus, under our assay condition, we reason that the ATP shifts the conformational equilibrium of AblFL to the open state, which is favored by imatinib over SKI binding.

### Inhibition of Abl kinase activity under double-drugging condition

The key question for clinical application is: What is the effect of different dosing concentration combinations of the two inhibitors on Abl’s kinase activity? Therefore, we performed synergy studies on AblFL kinase activity varying the concentration of both orthosteric and allosteric inhibitors **(Fig. 4*BC*)**. These experiments underscore the negative cooperativity between imatinib and asciminib, and corroborate the positive cooperativity between SKI and asciminib. First, we find a more pronounced inhibition of AblFL by SKI than by imatinib in the presence of asciminib. On the other hand, when used as a single agent, imatinib inhibits AblFL stronger than SKI, highlighting the difference in cooperativity. Second, we observe that in the presence of asciminib, less SKI is required for 90% inhibition of Abl activity compared to imatinib due to the positive cooperativity between SKI and asciminib **(Fig. 4*BC*)**.

### X-ray crystal structure of the ternary complex of AblFL-SKI-asciminib

Intrigued by the synergistic effect of SKI and asciminib on Abl activity, we structurally characterized this ternary complex by co-crystallization, resulting in a 2.86 Å crystal structure of AblFL-SKI-asciminib **(Fig. 5*A*, Fig. S12, Table S1)**. Surprisingly, this AblFL structure adopts a closed conformation with striking differences to previously reported closed structures; AblFL in complex with nilotinib and asciminib (PDB-ID: 5MO4), as well as in complex with PD166326 and myristic acid, the first groundbreaking structure of full-length Abl in the inhibited state (PDB-ID: 1OPK) **(Fig. 5*B*, Fig. S13)** (12, 35). First, we note that the entire N-terminal lobe is ~30° twisted only for AblFL-SKI-asciminib, when aligned by the regulatory domains **(Fig. 5*B*, Fig. S13)**. Second, the α-C helix is adopting the “out” position resulting from this N-lobe twist, since an α-C helix “in position” would clash with strands β4 and β5 **(Fig. 5*BC*)**. In consequence, the canonical salt-bridge between K290 and E305, a hallmark of an active kinase, is broken in our structure. Paradoxically, the two other closed ternary complexes of AblFL possess an α-C helix located in the “in position” and an established canonical salt-bridge (K290-E305), both reminiscent of an active kinase conformation **(Fig. S15)**. This highlights that, through binding of SKI and asciminib, we were able to capture the strictly inactive conformation of AblFL for the first time. We note, that the orthosteric site is fully occupied by SKI and the twisted N-lobe aids in forming this tightly packed binding pocket **(Fig. 5*C*, Fig. S13)**. In fact, K290 located on β3 strand is wrapping over SKI burying the inhibitor in AblFL’s orthosteric site. Besides extensive van-der-Waals interactions between SKI and AblFL, the quinazoline ring of SKI shares two hydrogen bonds with AblFL, one between the side chain hydroxyl of T334 on β-strand 5 and *N-2* of SKI as well as between the amide of M337 and *N*-0 of SKI **(Fig. 5C)**.

**Figure 5:**
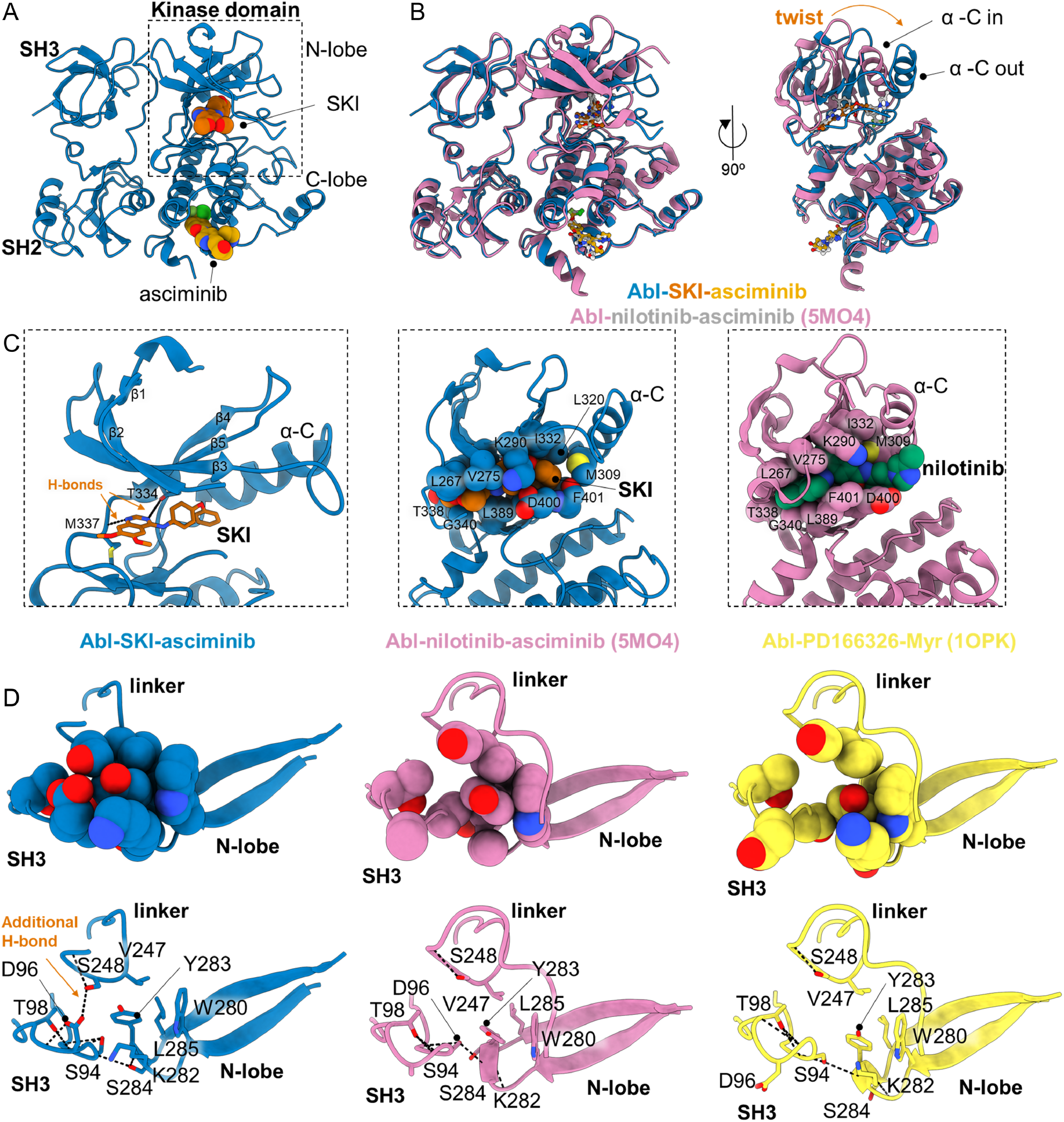
X-ray structure of ternary AblFL-SKI-asciminib complex reveals first fully-closed conformation compared to previous “energetically frustrated” ternary closed AblFL structures. (A) AblFL bound to SKI and asciminib. (B) Superposition of AblFL-SKI-asciminib (blue) and AblFL-nilotinib-asciminib (pink, PDB-ID: 5MO4) (12). When superimposed by the regulatory SH2 and SH3 domains, the N-lobe of AblFL-SKI-asciminib twists and exhibits α-C helix “out” position. (C) Zoom into the SKI and nilotinib binding sites. Van-der-Waals radii for the interacting Abl residues (spheres) with SKI (orange) and nilotinib (green) show more confined binding pocket for SKI than nilotinib. (D) Comparison of interface residues between N-SH3 domain (S94, D96, T98), linker (V247, S248), and N-lobe of kinase domain (W280, K282, Y283, S284, L285) for AblFL-SKI-asciminib (blue), AblFL-nilotinib-asciminib (pink, PDB-ID: 5MO4), and AblFL-PD166326-myristate (yellow, PDB-ID: 1OPK) (12, 35). Due to the twist in the N-lobe for AblFL-SKI-asciminib, residues in the domain/domain interface exhibit better packing. For AblFL-SKI-asciminib, additional hydrogen bond is established between S248 and D96 which contributes to this extended interface. Oxygen and nitrogen atoms are colored in red and blue, respectively. Carbon atoms are colored according to their respective protein cartoon.

When compared to other closed AblFL structures, we find an extended domain interface between SH3, linker and N-lobe of the kinase domain, which explains the positive cooperativity between SKI and asciminib. This improved interface is a direct result of the twisted N-lobe. The repositioned β2 and β3 strands cause Y283 to be completely buried within this interface. In other ternary complexes of AblFL, this interface is only partially formed **(Fig. 5*D***). Moreover, S248 (located on the linker) forms a hydrogen bond with D96 in the SH3 domain, which is only present in the ternary complex of AblFL-SKI-asciminib. Strikingly, S248P was identified as a resistant mutation for GNF-2 and asciminib in cell culture-based screening (7, 45) with no mechanistic understanding, given that this mutation is far away from the allosteric inhibitor binding site. Our structure now reveals the important role of S248 in allosteric closing of AblFL, and hence asciminib inhibition! We conclude that our complex of AblFL-SKI-asciminib represents the first example of a fully-closed and inactive Abl structure.

## Discussion

To combat on-target cancer drug resistance, double-drugging holds promise to be a powerful strategy. The rationale behind is multiplication of individual resistance mutational probabilities for each drug. Impressively, combinations of asciminib and various orthosteric inhibitors, including imatinib, indeed abolish the emergence of Abl resistance mutants and this double-drugging of Abl is currently in clinical trials (12). Given these groundbreaking clinical results, we used Abl kinase to interrogate the biophysical mechanism underlying this drug combination to learn a quantitative biophysical framework for successful double-drugging. In a second step, we used our knowledge of conformational equilibria in kinases to rationally select alternative orthosteric drugs exhibiting improved synergy with the allosteric drug. Our results have major implications: (i) Knowledge of conformational equilibria in drug targets indeed enables rational selection of inhibitor combinations with positive cooperativity and therefore better synergy. (ii) Our new fully-closed Abl structure solves the apparent mystery of all previous closed Abl structures with the α-C helix in the active “in position” that contradicted the common features of inactive-closed kinases with the α-C helix in the canonical “out position”. Structural investigation of our newly double-drugged ternary complex with SKI and asciminib reveals the first true α-C helix “out” state observed in Abl structures **(Fig. S14)**. This originates from a SKI-induced twist in the N-lobe causing a fully-closed conformation, thus, releasing an energetically frustrated conformation observed in other double-drugged AblFL complexes, since SKI and asciminib both preferentially bind to the closed state of AblFL to cause positive cooperativity. In contrast, double-drugging with an open conformation binder (nilotinib) and a closed conformation binder (asciminib) results in an energetically frustrated AblFL structure (PDB-ID: 5MO4). This structural study highlights how understanding of conformational equilibria crucially aids the discovery of further inhibited states. Our finding of the energetically frustrated conformation agrees with previous NMR experiments reporting an opposing binding preference of imatinib and GNF-5 for AblFL (32). In contrast, Johnson *et al*. claimed that such an antagonism arises from mutually exclusive binding of orthosteric and allosteric inhibitors (38). This conclusion contradicts previous studies characterizing AblFL-imatinib-GNF-5 by NMR as well as crystallographic studies on the ternary complex of AblFL bound to both nilotinib and asciminib (12, 32). Our ITC studies resolve this controversy by ruling out mutual exclusivity for binding of imatinib and asciminib. (iii) Our Abl data solve a heated debate: Recently, Kalodimos and colleagues argued that imatinib opens Abl via binding to its allosteric site (reported *K*_d_ >10 μM), and not via binding to its active site (46). This is in disagreement with NMR and cellular studies by Grzesiek *et al*. (32–34). Tighter binding of imatinib to open AblKD (*K*_d_ = 15 nM) compared to closed AblFL (*K*_d_ = 72.4 nM) and negative cooperativity between imatinib and asciminib buttress Grzesiek’s model where imatinib’s preferential binding to the open conformation of Abl arises from its orthosteric site binding with nanomolar affinity.

Double-drugging has been applied to two additional targets, SHP2 phosphatase (16) and EGFR kinase (15, 47). Fodor *et al*. used a combination of two allosteric binders, SHP099 and SHP504, to inhibit the phosphatase SHP2 (16).The authors demonstrate that the combination reduces the dosage requirements of these allosteric inhibitors to achieve effective inhibition of SHP2, however SHP504 is a very weak binder with an IC_50_ of 21 μM (16). For EGFR kinase, a combination of the inhibitor JBJ-04-125-02 binding right next to the irreversible orthosteric inhibitor osimertinib has been found to be more efficacious, than single agents, for inhibiting tumor growth in a mouse model. Furthermore, Jänne and colleagues demonstrated that this double-drugging resulted in the reduced emergence of resistance mutants in cellular assays (47). Here, the allosteric inhibitor binding site is in immediate proximity to the orthosteric site, resulting in direct interactions between the two inhibitors potentially driving positive cooperativity (15, 47).

In contrast, we investigated the mechanism of dual inhibition in AurA and Abl kinase targeting a distant allosteric site that is involved in natural regulation, in combination with active site drugs. Rationally targeting those natural allosteric sites has the advantage that it assures allosteric coupling to activity. We demonstrate with our first amateur attempts on both kinases that rational selection of double-drug combinations with positive cooperativity, and hence increased synergy, is possible based on knowledge of involved conformational equilibria.

In summary, this work proposes a biophysical framework for designing and evaluating double-drugging synergy utilizing orthosteric and allosteric modulators. As highlighted here, positive cooperativity is desirable for double-drugging approaches improving selectivity and dosage requirements. However, while extreme negative cooperativity is undesirable, the clinical success of Novartis’ drug combination for Abl (12) with 4-fold negative cooperativity as measured here suggests a clinical efficacy window ranging from small negative to strong positive cooperativity, given single-drug efficacy. Single drug efficacy is crucial, as otherwise a single resistance mutation abolishing binding of one drug would render the dual treatment to combat drug resistance essentially ineffective.

## Acknowledgements

D.K. is supported by the Howard Hughes Medical Institute (HHMI). The Berkeley Center for Structural Biology is supported in part by the Howard Hughes Medical Institute. The Advanced Light Source is a Department of Energy Office of Science User Facility under Contract No. DE-AC02-05CH11231. The Pilatus detector on 5.0.1. was funded under NIH grant S10OD021832. The ALS-ENABLE beamlines are supported in part by the National Institutes of Health, National Institute of General Medical Sciences, grant P30 GM124169.

## Author Contributions

C. K., A.H., V.N. and D.K. conceived the project and designed the experiments. C.K. performed all biochemical experiments and analyzed the data. V.N. purified AurA and Mb1. V.N. also performed the initial ITC experiments. A.H. and S.K. solved AurA-danusertib-Mb2 and AurA-danusertib-Mb1 crystal structures, respectively. H.L. further modeled, refined and analyzed AurA-Mb1-danusertib and AurA-Mb2-danusertib structures. C.K. and H.L. crystalized the AblFL-SKI-asciminib complex. H.L obtained and processed X-ray diffraction data. C.K. and H.L solved, modeled, refined and analyzed this structure. C.K., H. L. and D.K. wrote the paper. All authors commented on the manuscript and contributed to data interpretation.

## Methods

### Cloning and Purification of Aurora A and monobodies

Dephosphorylated AurA (residues 122-403, TEV-cleavable, N-terminal His6-tagged, kanamycin-resistance) in pET28a and LPP (#79748) from Addgene were cotransformed in BL21(DE3) cell and plated on Kan/Spec LB plate. Few robust colonies were grown overnight in 250 ml TB culture to inoculate a 1 L culture to OD = 0.2. 1 L TB culture was grown to OD = 0.6 – 0.8 and induced with 0.6 mM IPTG for 16 hours at 21 °C. Cells were centrifuged at 4500 rpm for 15 min and resuspended in Buffer A. Cells were sonicated in the presence of EDTA-free protease inhibitor cocktail, lysozyme and DNAse, and then spun down at 18000 rpm. Supernatant was filtered through 0.22 μm filtering unit and passed through Ni-NTA columns. The protein was eluted at 100% Buffer B and protein fractions were pooled and combined with TEV and GST-LPP, then dialyzed overnight against Buffer C at 4 °C. Cleaved Aurora A was passed through tandem Ni-NTA and GST columns to remove uncleaved Aurora A, cleaved His-tag, and GST-LPP, and ran through 26/600 S200 pg gel filtration column with Buffer D. Pure fractions were pooled and concentrated to around 40 μM, and stored in −80 °C.

Buffer A: 50 mM Tris-HCl, 300 mM NaCl, 20 mM MgCl_2_, 10% glycerol, pH 8.0 Buffer B: 50 mM Tris-HCl, 300 mM NaCl, 500 mM imidazole, 20 mM MgCl_2_, 10% glycerol, pH 8.0 Buffer C: 50 mM Tris-HCl, 300 mM NaCl, 1 mM MnCl_2_, 5 mM TCEP, 10% glycerol, pH 7.5 Buffer D: 20 mM Tris-HCl, 200 mM NaCl, 20 mM MgCl_2_, 5 mM TCEP, 10% glycerol, pH 7.5 Monobodies (TEV-cleavable, N-terminal His6-tagged) were purified with on-column refolding as described in Zorba *et al*. (25).

### Cloning and Purification of AblKD and AblFL

Plasmids (kanamycin-resistance) encoding AblKD (residues 229-510, TEV-cleavable, N-terminal MBP-His6-tagged) and AblFL (residues 64-510, TEV-cleavable, N-terminal MBP-His6-tagged) was synthesized and cloned by GenScript into pETm41 **(Fig. S7)**. Residue numbering follows Abl1b isoform that naturally consists of N-myristoylation. All Abl constructs were co-expressed with phosphatase YOPH (streptomycin-resistance) in BL21(DE3) cells and plated on Kan/Strep LB plate. Few robust colonies were grown overnight in 250 ml TB culture to inoculate a 1 L culture to OD = 0.2. Six of 1 L TB cultures were grown to OD = 0.6 – 0.8 and induced with 0.1 mM (for AblKD) or 0.2 mM IPTG (for AblFL) for 16 – 20 hours at 18 °C. Cells were centrifuged at 4500 rpm for 15 mins and resuspended in Buffer E. Cells were sonicated in the presence of EDTA-free protease inhibitor cocktail, lysozyme and DNAse, and then spun down at 18000 rpm. Supernatant was filtered through 0.22 μm filtering unit and passed through Ni-NTA columns. The protein was eluted at 100% Buffer F and protein fractions were pooled and combined with TEV and CIP (#M0525) from NEB and dialyzed overnight against Buffer E at 4 °C. Cleaved Abl was passed through Ni-NTA column to get rid of uncleaved Abl, cleaved His-tag. Cleaved Abl fractions were pooled and dialyzed against Buffer G, and ran through Q-column to remove remaining phosphorylated Abl and impurities with gradual elution using Buffer H. Dephosphorylated Abl fractions were ran through 26/600 S75 pg (for AblKD) or 26/600 S200 pg (for AblFL) gel filtration column with Buffer E. Pure fraction was aliquoted to around 40 μM, and stored in −80 °C.

Buffer E: 50 mM Tris-HCl, 500 mM NaCl, 1 mM TCEP, pH 8.0 Buffer F: 50 mM Tris-HCl, 500 mM NaCl, 500 mM imidazole, 1 mM TCEP, pH 8.0 Buffer G: 50 mM Tris-HCl, 1 mM TCEP, 10% glycerol, pH 8.0 Buffer H: 50 mM Tris-HCl, 1 M NaCl, 1 mM TCEP, pH 8.0

### Isothermal titration calorimetry

All titrations were carried out using Nano ITC (TA Instruments) and analyzed via the NanoAnalyze software either using the independent fit model or competitive replacement model. The first injection of each experiment was discarded according to the software manual.

For AurA, danusertib (Selleckchem #S1107) was reconstituted to 100 mM in 100% DMSO and were diluted to appropriate concentration to match final 5% DMSO (vol/vol) for each experiment. An ADP-analogue, AMPCP, was used for competitive replacement experiments to measure and fitting of the binding of danusertib to AurA. All proteins were dialyzed in 20 mM Tris-HCl, 200 mM NaCl, 10% (vol/vol) glycerol, 5 mM TCEP, pH 7.5. AMPCP was resuspended with the same buffer and was matched to pH 7.5. DMSO was added prior to each experiment to match 5% between titrant and titrand. Each injection was added in 2 μl increments with 180 s interval at a constant stirring speed of 300 rpm and at 25°C. Concentrations used for the experiments are noted below.

20 μM AurA + 150 μM Mb1
20 μM AurA / 50 μM Danusertib + 384 μM Mb1
20 μM AurA / 150 μM Danusertib + 384 μM Mb1
20 μM AurA + 150 μM Mb2
20 μM AurA / 150 μM Danusertib + 150 μM Mb2
20 μM AurA + 90.9 μM Mb3
20 μM AurA / 150 μM Danusertib + 90.9 μM Mb3
100 μM AurA + 1290 μM AMPCP
20 μM AurA / 2000 μM AMPCP + 150 μM Danusertib
40 μM AurA / 140 μM Mb1 + 1000 μM AMPCP
20 μM AurA / 2000 μM AMPCP / 140 μM Mb1 + 150 μM Danusertib
80 μM AurA / 200 μM Mb2 + 977.6 μM AMPCP
20 μM AurA / 2000 μM AMPCP / 200 μM Mb2 + 150 μM Danusertib
80 μM AurA / 100 μM Mb3 + 1200 μM AMPCP
20 μM AurA / 2000 μM AMPCP / 100 μM Mb3 + 150 μM Danusertib
8 μM AurA + 51.7 μM Mb6
8 μM AurA + 25 μM Danusertib + 51.7 μM Mb6

For Abl, N-Myr peptide (Myr-GQQPGKVLGDQR), ordered from Genscript, was used for competitive replacement experiments to measure and fitting of the binding of asciminib to Abl. All proteins were dialyzed in 50 mM Tris-HCl, 500 mM NaCl, 1 mM TCEP, pH 8.0. Imatinib-mesylate (Sigma #SML-1027) and asciminib (MedKoo #206490) were reconstituted to 10 mM in 100% DMSO and were diluted to appropriate concentration for each experiment. N-Myr peptide was resuspended with the same buffer and was matched to pH 8.0. DMSO was added prior to each experiment to match 5% between titrant and titrand. Each injection was added in 1-1.5 μl increments with 180 s interval at a constant stirring speed of 300 rpm and at 25°C. Concentrations used for the experiments are noted below.

15 μM AblKD + 100 μM Imatinib
15 μM AblKD / 50 μM Asciminib + 100 μM Imatinib
15 μM AblFL + 100 μM Imatinib
15 μM AblFL / 50 μM Asciminib + 100 μM Imatinib
15 μM AblFL / 150 μM Asciminib + 100 μM Imatinib
30 μM AblKD + 400 μM N-Myr
30 μM AblKD / 50 μM Imatinib + 400 μM N-Myr
30 μM AblFL + 400 μM N-Myr
30 μM AblFL / 50 μM Imatinib + 400 μM N-Myr
25 μM AblKD / 300 μM N-Myr + 250 μM Asciminib
25 μM AblKD / 50 μM Imatinib / 300 μM N-Myr + 250 μM Asciminib
25.5 μM AblFL / 300 μM N-Myr + 250 μM Asciminib
25.5 μM AblFL / 50 μM Imatinib / 300 μM N-Myr + 250 μM Asciminib
30 μM AblKD + 2500 μM AMPPCP
30 μM AblFL + 2500 μM AMPPCP

### *In vitro* kinase assay

To measure the IC_50_ of danusertib to AurA, ADP-Glo^TM^ Max assay (Promega #V7001) was used. 20 nM AurA in the absence or presence of either saturating concentration of Mb1 or Mb2 or Mb3 was incubated with 3 mM Lats2 (ATLARRD**<u>S</u>**LQKPGLE), 0.6 mg/ml BSA and varying concentrations of danusertib with final 5% (vol/vol) of DMSO at 25 °C in 20 mM Tris-HCl, 200 mM NaCl, 10% (vol/vol) glycerol, 5 mM TCEP, pH 7.50. The bolded and underlined residue indicate site of phosphorylation. The reaction was initiated by adding 5 mM ATP, and the final samples were collected after 2 h for AurA-Mb1 complex, 10 h for apo AurA, and 20 h for AurA-Mb2 and AurA-Mb3 complexes. The amount of ADP in the samples were measured by following the manufacturer’s protocol and used to calculate the observed rate.

Assays for Abl were performed at 25 °C with half-well 96-well plate (Corning #3994) in 50 mM Tris-HCl, 500 mM NaCl, 1 mM TCEP, pH 8.0 supplemented with 20 nM Abl kinase (AblKD or AblFL), 2 mM Srctide (EI**<u>Y</u>**GEFKK), 0.6 mg/ml BSA, 20 mM MgCl_2_, 750 μM NADH, 6 mM PEP, 2.5 units of PK/LDH (Sigma #P0294). The bolded and underlined residue indicate site of phosphorylation. Oxidation of NADH at A_340_ was monitored using SpectraMAX by starting the assay with 1 mM ATP. Final volume of the assay was 100 μL. Observed rate (*k*_obs_) was calculated following Zorba *et al*. (25). All data were processed using GraphPad Prism and fitted to a four-parameter doseresponse model.

### Molecular dynamics simulation

All-atom molecular dynamics simulations were conducted using OpenMM 7.6 (48) and ‘Making it rain’ cloud-based notebook environment (49). The structure of AurA-Mb1-danusertib was used as an initial model. To mimic danusertib binding to AurA-Mb1 under ITC conditions, we created such structure via removal of danusertib from our ternary complex AurA-danusertib-Mb1 (since the published AurA-Mb1 structure (PDB-ID: 5G15) has AMPPCP bound to active site (25). Parameterization for all MD runs was conducted using LEaP (50) with Amber ff14SB force field (51), GAFF2 (52) for ligand and TIP3P (53, 54) water model. The systems were neutralized with NaCl at 0.2 mM, following the ITC conditions, and box size was set at 20 Å. AurA-Mb1 and AurA-Mb1-danusertib structures were equilibrated to 298 K via Langevin dynamics (55) and 1 bar via Monte Carlo barostat (56) with 2 fs integration time. We set 10,000 steps of energy minimization with 1,000 kJ/mol of harmonic position restraints. The systems were equilibrated for 0.2 ns and 1 ns for AurA-Mb1 and AurA-Mb1-danusertib, respectively, in the NVT ensemble. Then, with accordingly equilibrated systems, triplicates of 10 ns production runs were done in the NPT ensemble. Trajectories were analyzed using VMD 1.9.4a53 (57).

### FRET measurements

FluoroMax-4 (Horiba Scientific) with temperature controller (water bath) was used to measure FRET between intrinsic tryptophan fluorescence and SKI. Either 10 nM Abl or 10 nM Abl + 200 nM asciminib were pre-incubated with varying concentrations of SKI for 40 min at 25 °C before measurements. An increase in the fluorescence was measured when the complex, specifically tryptophan, was excited at 295 nm to emit at 340 nm, which then excites SKI to emit at 460 nm **(Fig. S9)**. Both 5 nm of excitation and emission slit width were used. Control experiments (buffer-only, protein-only, and inhibitor-only) were confirmed that the increase of fluorescence is caused by the fluorescence energy transfer.

The fluorescence intensity at 460 nm versus SKI concentration was fitted to quadratic equation below in GraphPad Prism to obtain apparent *K*_d_.

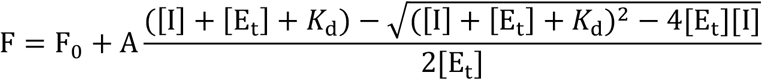

We simulated curves with tighter *K*_d_ for comparison to ensure that the fitted curves are not step-functions due to the high enzyme concentration **(Fig. S10)**.

### Crystallographic methods

Crystals of AurA in complex with Mb1 and danusertib were obtained by combining 2 μl of 300 μM (10 mg/mL) AurA + 315 μM (4 mg/mL) Mb1 + 2 mM AMPPCP + 4 mM MgCl_2_ with 2 μl reservoir of 0.1 M MES pH 6.5 + 0.2 M Ammonium sulfate + 4% (v/v) 1,3-propanediol + 15-18% PEG 8000. Streak seeding was used to obtain bigger crystals. Crystals were grown at 18 °C by hanging drop. The crystals were transferred to a drop of fresh reservoir for 30 seconds to remove excess of nucleotides from the crystal surface. Then, the crystals were transferred to a drop with reservoir with 1 mM danusertib for 16 h of soaking. For cryo-protection, the crystals were transferred into 17.5% PEG 400, 17.5% ethylene glycol, 15% reservoir, 50% water for a few seconds.

Crystals of AurA in complex with Mb2 and danusertib were obtained by combining 0.5 μl of 300 μM (10 mg/mL) AurA + 315 μM (4 mg/mL) Mb2 + 1 mM danusertib with 0.5 μl of 0.1 M BIS-TRIS pH 5.5 + 0.2 M Ammonium acetate + 25% PEG3350. Crystals were grown at 18 °C by sitting drop. Crystals were harvested and subsequently flash frozen. Diffraction data for AurA-danusertib-Mb1 and AurA-danusertib-Mb2 were collected at 100 K Advanced Light Source (Lawrence Berkeley National Laboratory) at beamlines BL821 and BL501, respectively, and were integrated with XIA2 (58) or XDS (59). Data were scaled and merged with AIMLESS (60). Initial phases were obtained with molecular replacement programs MOLREP (61) and PHASER (62) by using AurA + Mb1 + AMPPCP (PDB-ID: 5G15) for AurA-danusertib-Mb1 structure, and AurA + AMPPCP (PDB-ID: 4C3R) and HA4Mb (PDB-ID: 3K2M) for AurA-danusertib-Mb2 structure using 2 molecules each in the asymmetric unit. The structures were iteratively refined using refmac and phenix.refine (Version1.19.1) (63) followed by manual model building in COOT (64). Models were validated with MolProbity (65). Molecular structures were represented and rendered with ChimeraX (66, 67).

Crystals of AblFL in complex with SKI and asciminib were obtained by combining 0.3 μl of 600 μM AblFL + 700 μM SKI + 700 μM asciminib (~32 mg/ml) in 5% DMSO with 0.4 μl reservoir of 0.1 M Tris-HCl pH 8 + 1.75 M Ammonium sulfate + 2% (v/v) polypropylene glycol 400 (PPG 400). The final stock of complex was concentrated from 1 μM AblFL with ~1.2 μM SKI/asciminib after incubation at 4 °C for 6 h. Screening around this condition yielded crystals in a transparent diamond-shaped or plate-shaped crystals. Crystals were grown at 18 °C by sitting drop for a few days. The crystals were transferred to a drop of fresh reservoir containing 20% Xylitol with matching concentration of inhibitors in 5% DMSO for few seconds for cryo-protection.

Single crystal X-ray diffraction data was collected at 100 K at Advanced Light Source (ALS) Berkeley (BL201). Data were integrated with XDS (59) as well as scaled and merged with AIMLESS (60). Analysis of processed data with phenix.xtriage (68) revealed substantial translational non-crystallographic symmetry (tNCS) with a Patterson peak of 56.63% height relative to origin. Initial phases were obtained by molecular replacement (PHASER) (62) using AblFL-nilotinib-asciminib (PDB-ID: 5MO4) as a search model with two molecules in the asymmetric unit. The kinase domain, SH3 and SH2 (regulatory domains) were individually placed during molecular replacement. Refinement and manual model building were performed by phenix.refine (version 1.19.1) and Coot, respectively (63, 64). Models were validated with MolProbity (65). Molecular structures were represented and rendered with ChimeraX (66, 67) and PyMol (69).

