## Supplemental Information for "A biophysical framework for double-drugging kinases"

Chansik Kim, Hannes Ludewig, Adelajda Hadzipasic, Steffen Kutter, Vy Nguyen, Dorothee Kern[1]

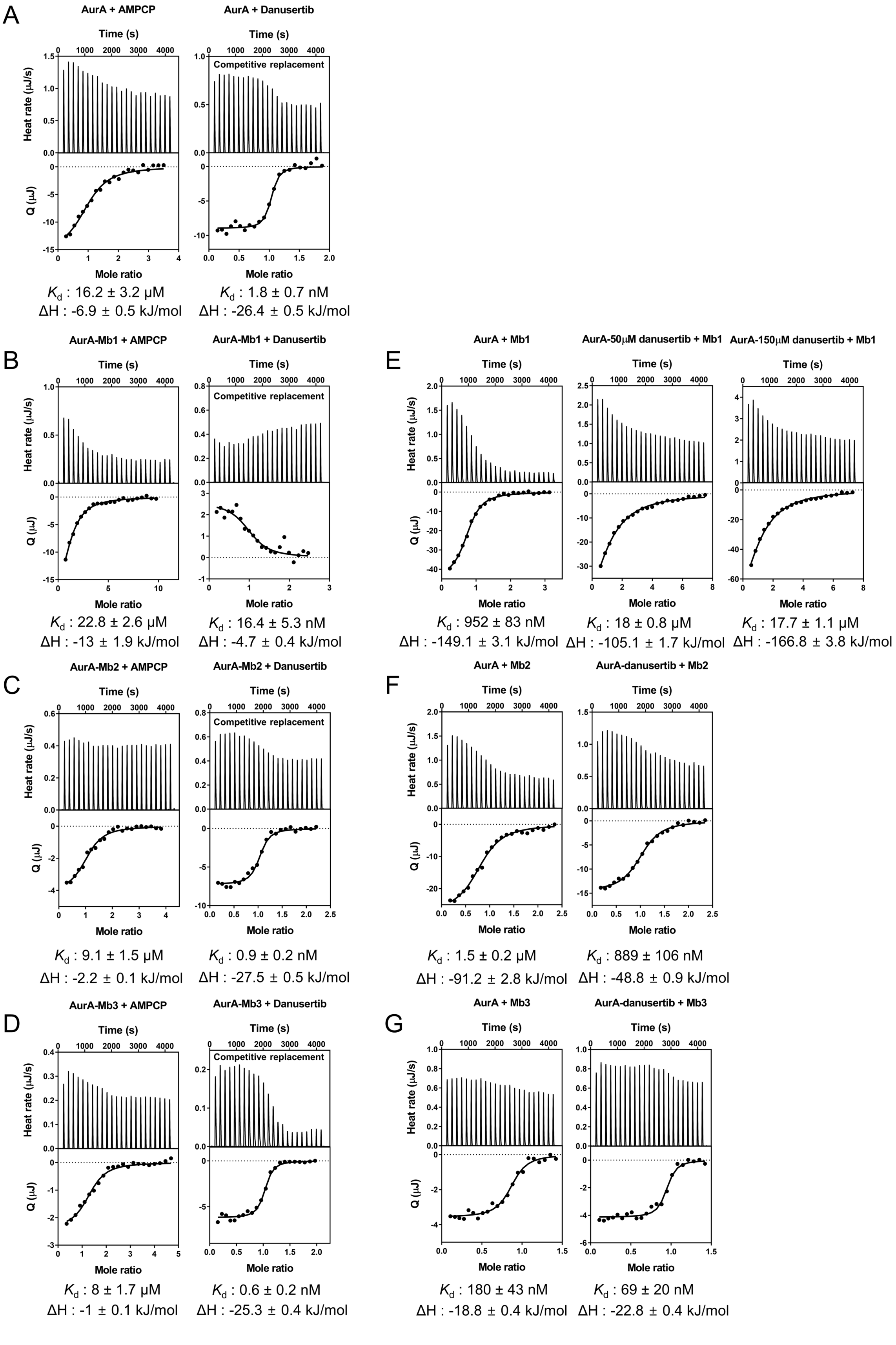
A
AurA + AMPCP
AurA + Danusertib
Competitive replacement
Time (s)
Heat rate (µJ/s)
Q (µJ)
Mole ratio
$K_d$ : 16.2 ± 3.2 µM
ΔH : -6.9 ± 0.5 kJ/mol
$K_d$ : 1.8 ± 0.7 nM
ΔH : -26.4 ± 0.5 kJ/mol
B
AurA-Mb1 + AMPCP
AurA-Mb1 + Danusertib
$K_d$ : 22.8 ± 2.6 µM
ΔH : -13 ± 1.9 kJ/mol
$K_d$ : 16.4 ± 5.3 nM
ΔH : -4.7 ± 0.4 kJ/mol
E
AurA + Mb1
AurA-50µM danusertib + Mb1
AurA-150µM danusertib + Mb1
$K_d$ : 952 ± 83 nM
ΔH : -149.1 ± 3.1 kJ/mol
$K_d$ : 18 ± 0.8 µM
ΔH : -105.1 ± 1.7 kJ/mol
$K_d$ : 17.7 ± 1.1 µM
ΔH : -166.8 ± 3.8 kJ/mol
C
AurA-Mb2 + AMPCP
AurA-Mb2 + Danusertib
$K_d$ : 9.1 ± 1.5 µM
ΔH : -2.2 ± 0.1 kJ/mol
$K_d$ : 0.9 ± 0.2 nM
ΔH : -27.5 ± 0.5 kJ/mol
F
AurA + Mb2
AurA-danusertib + Mb2
$K_d$ : 1.5 ± 0.2 µM
ΔH : -91.2 ± 2.8 kJ/mol
$K_d$ : 889 ± 106 nM
ΔH : -48.8 ± 0.9 kJ/mol
D
AurA-Mb3 + AMPCP
AurA-Mb3 + Danusertib
$K_d$ : 8 ± 1.7 µM
ΔH : -1 ± 0.1 kJ/mol
$K_d$ : 0.6 ± 0.2 nM
ΔH : -25.3 ± 0.4 kJ/mol
G
AurA + Mb3
AurA-danusertib + Mb3
$K_d$ : 180 ± 43 nM
ΔH : -18.8 ± 0.4 kJ/mol
$K_d$ : 69 ± 20 nM
ΔH : -22.8 ± 0.4 kJ/mol

**Figure S1: ITC profiles of AMPCP and danusertib binding to AurA and AurA-Mb complexes, and binding of Mbs to AurA and AurA-danusertib.** Apparent dissociation constants, $K_d$, and change in enthalpy, $\Delta H$, are shown with errors that represent 68.3% confidence interval of the fit of the data (± 1 s.d.). Binding of danusertib data were fitted with a competitive replacement model accounting for the $K_d$ of AMPCP binding. (A-D) Binding of danusertib to AurA is weakened by 18-fold in the presence of Mb1 due to Mb1-induced conformational equilibrium shift to the active state. In contrast, Mb2 and Mb3 shift the equilibrium to the inactive state so that the binding of danusertib is tightened by 2-fold and 3-fold, respectively. (E-G) Binding of monobodies to AurA are affected by identical fold-change due to pre-incubation of danusertib. For Mb1, two saturating concentrations of danusertib, 50 μM and 150 μM, were used during pre-incubation to confirm simultaneous binding of the opposite conformation binders, danusertib and Mb1, is possible.

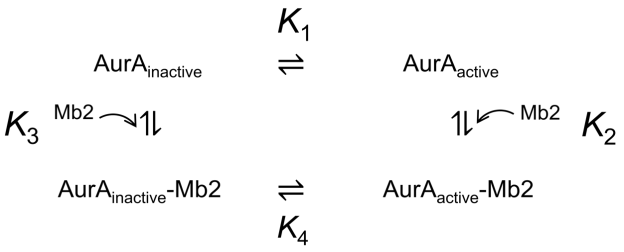


$$(1)\ K_{d\_Mb2\_obs} = \frac{([Aur_{act}] + [Aur_{inact}])[Mb2]}{([Aur_{act}{:}Mb2] + [Aur_{inact}{:}Mb2])} = \frac{K_3(K_1 + 1)}{K_4 + 1}$$

(2) $K_1 = 0.67 \pm 0.09$

(3) Danusertib fully shifts conformational equilibrium to inactive.
Then, $K_3 = K_d$ from AurA-danusertib + Mb2

$$(4)\ Fold\ change = \frac{K_{d\_Mb2\_obs}}{K_3} = \frac{(K_1 + 1)}{K_4 + 1} = \frac{1 + 0.67}{K_4 + 1}$$ ··· (Equation in (1))

(5) Mb2 fully shifts conformational equilibrium to inactive
Then, $K_4 \rightarrow 0$

$$(6)\ \therefore Fold\ change = \frac{1 + 0.67}{\cancel{K_4} + 1} = \frac{1 + 0.67}{1} = 1.67$$

∴ Theoretical maximum fold-change solely by shifting the active/inactive conformational equilibrium of AurA is 1.67-fold ± 0.09.

**Figure S2: Calculated maximal cooperativity for double-drugging by active/inactive equilibrium shift of AurA.** Apparent $K_d$ (Equation in (1)) for Mb2 binding can be derived from a reversible two-state allosteric model. By using the active/inactive equilibrium constant ($K_1$) from Warintra *et al.* (1) and assumption (3), we can simplify the equation to derive a fold-change in the apparent Kd of Mb2 after preincubation with danusertib (4). Since Mb2 binding fully shifts the conformational equilibrium of AurA to inactive ($K_4 \rightarrow 0$), maximal fold change is calculated to be 1.67 ± 0.09. Therefore, we reason, within experimental error, the 2-fold positive cooperativity between danusertib and Mb2 to AurA can be solely explained by the shift of the conformational equilibrium.

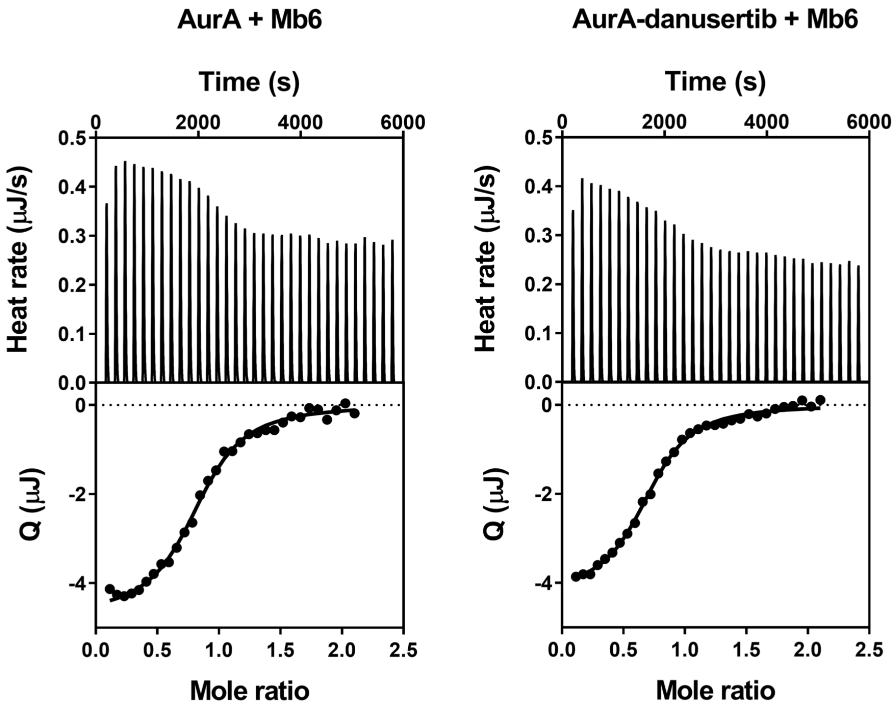


$K_d$ : 383 ± 33 nM
ΔH : -61.1 ± 0.98 kJ/mol

$K_d$ : 392 ± 30 nM
ΔH : -54.8 ± 0.81 kJ/mol

**Figure S3: ITC profiles of Mb6 binding to AurA and AurA-danusertib reveal identical affinities.** Zorba *et al.* discovered that Mb6 neither activates nor inhibits AurA despite tight binding to the allosteric site of AurA, implying that Mb6 does not shift of the active/inactive conformational equilibrium of AurA (2). Thus, we hypothesized, and then show here, that the binding of Mb6 is not affected by pre-incubation of danusertib, confirming the classic allosteric binding model between distant binding sites of AurA. Observed dissociation constants, $K_d$, and change in enthalpy, $\Delta H$, are shown with errors that represent 68.3% confidence interval of the fit of the data (± 1 s.d.).

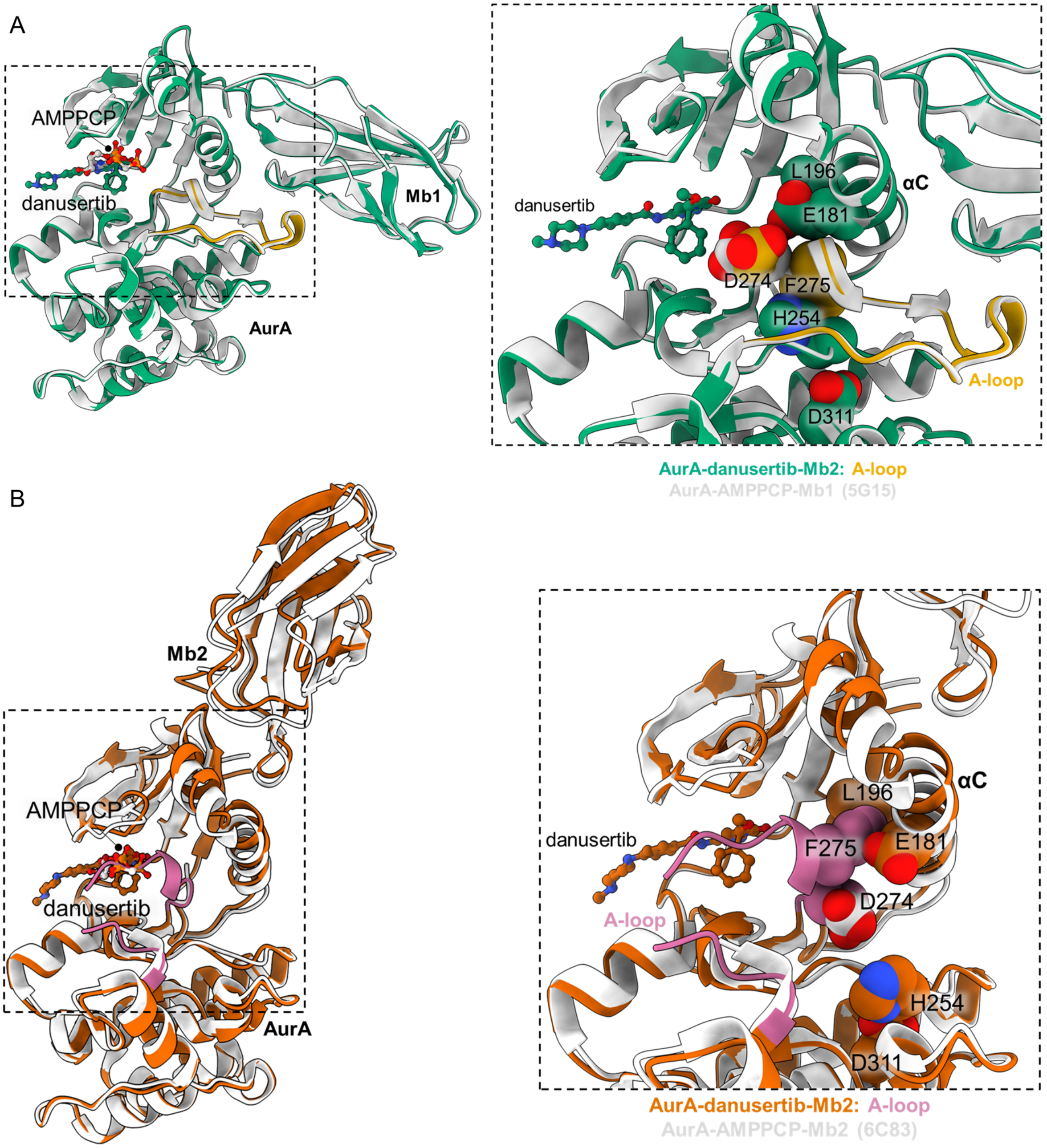


**Figure S4: Comparison of ternary complex X-ray structures of AurA-danusertib-Mb1 and AurA-danusertib-Mb2 to its binary counterparts AurA-Mb1 and AurA-Mb2.** (A) Superposition of AurA-danusertib-Mb1 (green) and AurA-AMPPCP-Mb1 (white, PDB-ID: 5G15) show that the hallmarks of an active kinase are still present in AurA-danusertib-Mb1 complex (extended A-loop, gold, intact regulatory spine, shown as spheres right panel). However, the active site D274 is rotated away from the terminal phenyl ring of danusertib to avoid a steric clash. We note that the AurA-danusertib-Mb1 structure was obtained by soaking AurA-AMPPCP-Mb1 crystals with danusertib replacing AMPPCP. (B) Superposition of AurA-danusertib-Mb2 (orange) and AurA-AMPPCP-Mb2 (white, PDB-ID: 6C83) show all hallmarks of an inactive kinase (A-loop folded towards the active site and partially missing electron density, regulatory spine broken, shown as spheres).

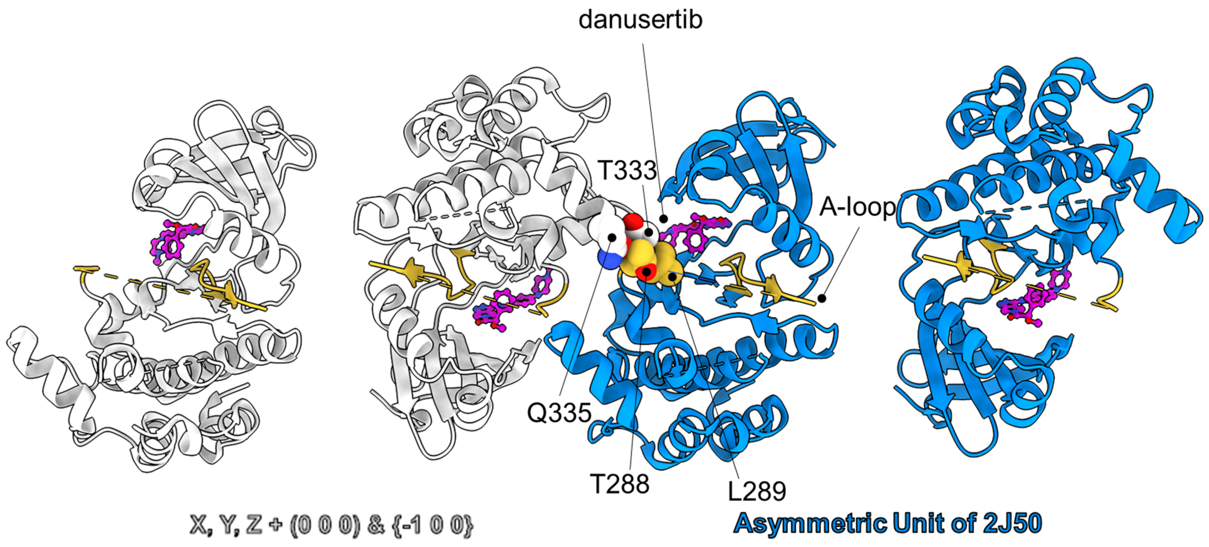


**Figure S5: Crystal packing contacts in AurA-danusertib (PDB-ID: 2J50) (3) as possible reason for activation loop conformation.** Our x-ray structure of the ternary complex AurA-danusertib-Mb2 had all hallmarks of the inactive kinase conformation **(Fig. 2*B*)**. The published X-ray structure of the AurA-danusertib complex shows AurA in the inactive conformation with respect to the regulatory spine, but not the activation loop. The activation loop inactive conformation may not be observed in 2J50 due to crystal contacts between residues (spheres) on the activation loop (yellow) in one monomer in the asymmetric unit (blue), and residues (white, sphere) of another monomer from the -1 0 0 symmetry mate (white). Danusertib is shown in sphere-and-stick representation in magenta. Oxygen and nitrogen atoms are colored in red and blue, respectively. Carbon atoms are colored according to their respective protein cartoon. Dotted lines represent unresolved parts of the structure (residues 280-287, 303-306).

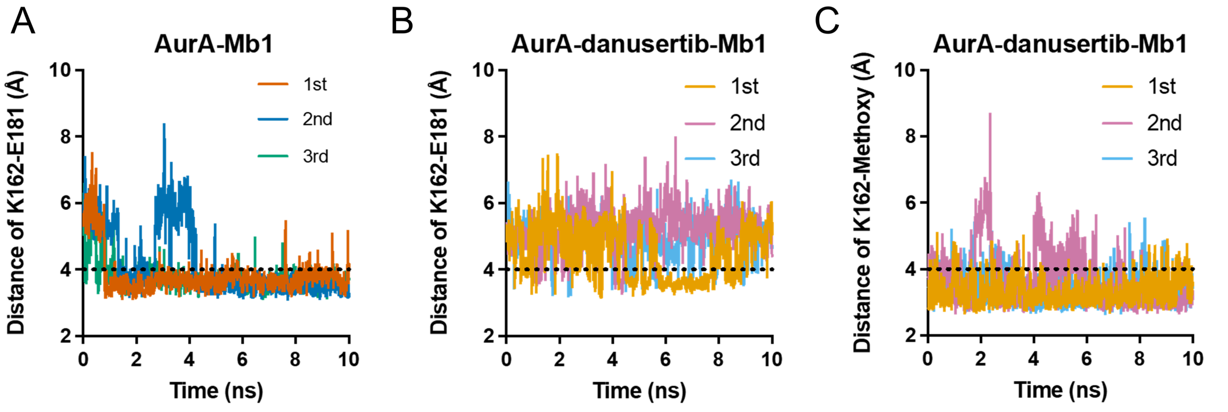


1st: 84.8%, 2nd: 70.2%, 3rd: 87.5%    1st: 38.9%, 2nd: 5.6%, 3rd: 7.3%    1st: 89.8%, 2nd: 69.7%, 3rd: 83.7%

**Figure S6: Molecular dynamics simulation of AurA to study the effect of danusertip binding on the K162 - E181 salt bridge in AurA and the H-bond bewteen *O*-27 (methoxy) of danusertib and K162 (See also Fig. 2*C-H*).** Three independent 10 ns MD runs ($1^{st}$, $2^{nd}$, $3^{rd}$) were performed for each complex (see Methods for details). (A) The K162-E181 salt-bridge is established, indicated by high salt-bridge occupancy, in MD simulations when danusertib is removed from AurA-Mb1-danusertib structure. Distance cut-off for salt-bridge is set to lower than 4 Å (dashed line) (B) When danusertib is not removed, K162-E181 is rarely established, (C) and K162 (*N*-ζ) interacts with *O*-27 of methoxy moiety of danusertib instead. Distance cut-off for salt-bridge and hydrogen bond are set to lower than 4 Å (dashed line).

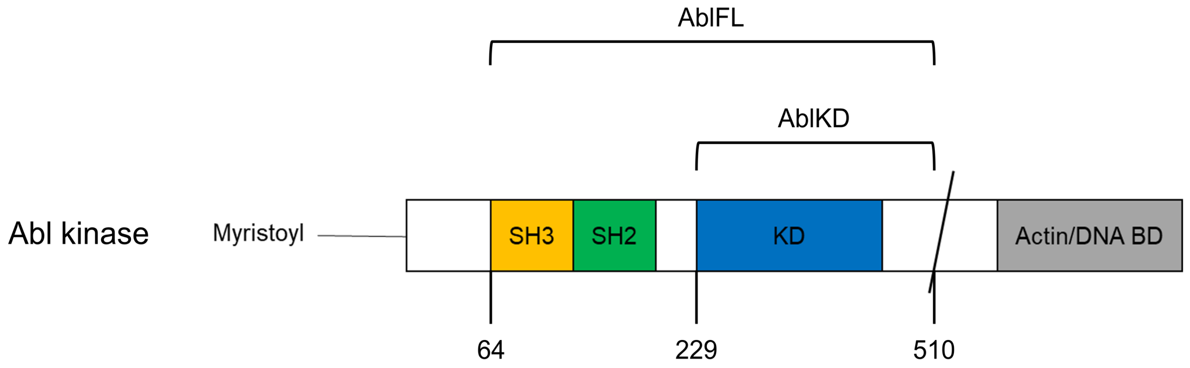


**Figure S7: Defining the Abl constructs used in this study.** AblFL (64-510) includes SH3, SH2 regulatory domains and kinase domain (KD). AblKD (229-510) only consists of KD.

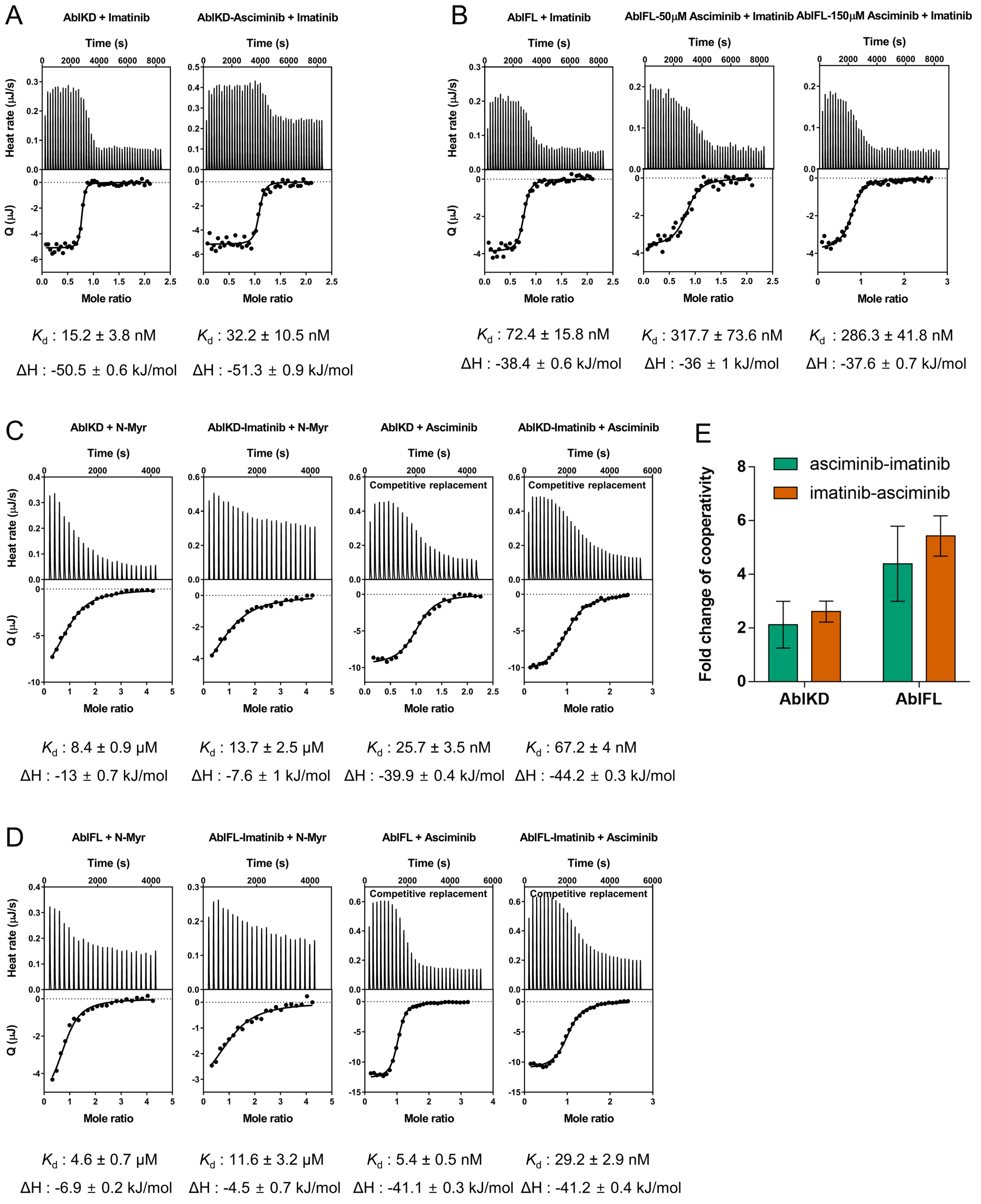


**Figure S8: ITC profiles of imatinib binding to AblKD and AblFL with/without asciminib and N-Myr peptide as well as asciminib binding to AblKD and AblFL with/without imatinib.** Observed dissociation constants, $K_d$, and change in enthalpy, Δ$H$, are shown with errors that represent 68.3% confidence interval of the fit of the data (± 1 s.d.). Binding of asciminib data were fitted with a competitive replacement model accounting for the $K_d$ of N-Myr peptide binding. (A-B) Imatinib binds tighter to AblKD than to AblFL due to its preference for the open form of Abl. Pre-incubation of asciminib to AblKD and AblFL weakens the binding affinity of imatinib by 2-fold and 4-fold, respectively. Two concentrations (50 μM and 150 μM) of asciminib were used during pre-incubation to confirm the simultaneous binding of imatinib and asciminib to AblFL. (C-D) Binding of asciminib to AblKD and AblFL were measured using competitive replacement ITC with N-Myr, a weak allosteric binder. Identical fold of negative cooperativity was found, within error (see (E)), regardless of asciminib and imatinib binding order. We find comparable binding affinities of N-

Myr or asciminib to AblFL-imatinib and to AblKD, indicating imatinib shifts the open/closed conformational equilibrium of Abl to the open via binding to orthosteric site. In this experimental set-up, imatinib cannot occupy the allosteric site due to the presence of N-Myr. (E) Within the range of error, fold change in cooperativities between imatinib and asciminib are identical regardless of binding order. Errors represent propagated errors from ITC data.

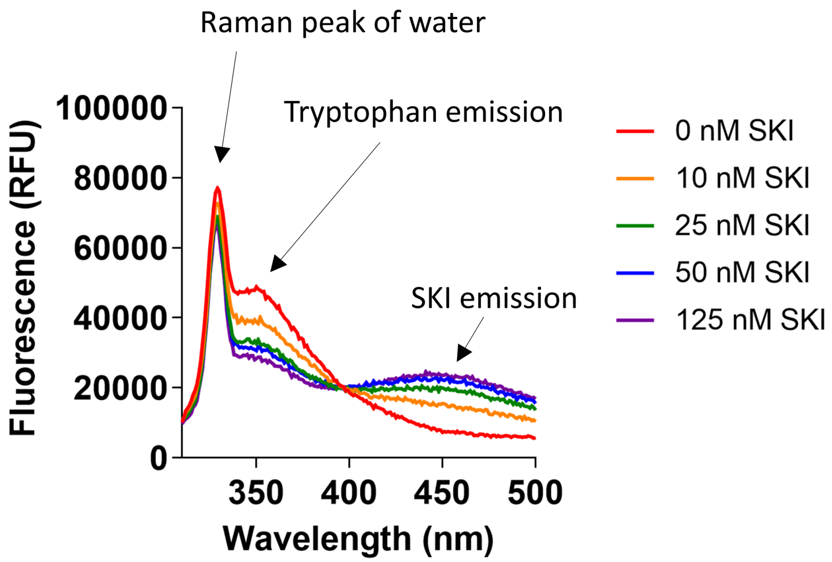


**Figure S9: FRET experiments to detect SKI binding at 25 ºC with 25 nM AblFL.** Excitation of tryptophan at 295 nm emits at around 350 nm, where SKI is excited and emits at 460 nm to observe the binding of SKI to Abl. Decrease in tryptophan emission at 350 nm is directly proportional to increase in SKI emission at 460 nm.

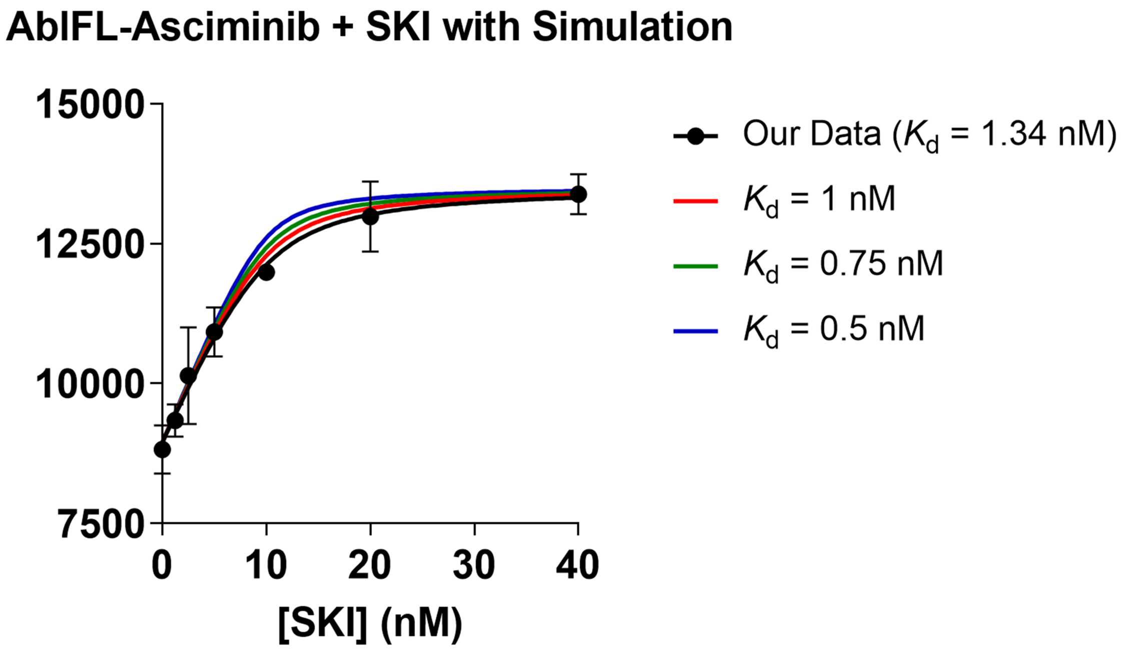


**Figure S10: Simulated SKI binding curves for tighter $K_d$ in FRET data.** We fit our data (*n* = 3, mean ± s.d.m.) to a quadratic binding equation (see Methods for details) due to lower $K_d$ than AblFL concentration used (10 nM). The simulation of binding curves with $K_d$ of 1 nM (red), 0.75 nM (green), 0.5 nM (blue) would show steeper increase in signal than our experimental data, indicating that our experimental data is not yet yielding a step-function and the fitted $K_d$ can be trusted.

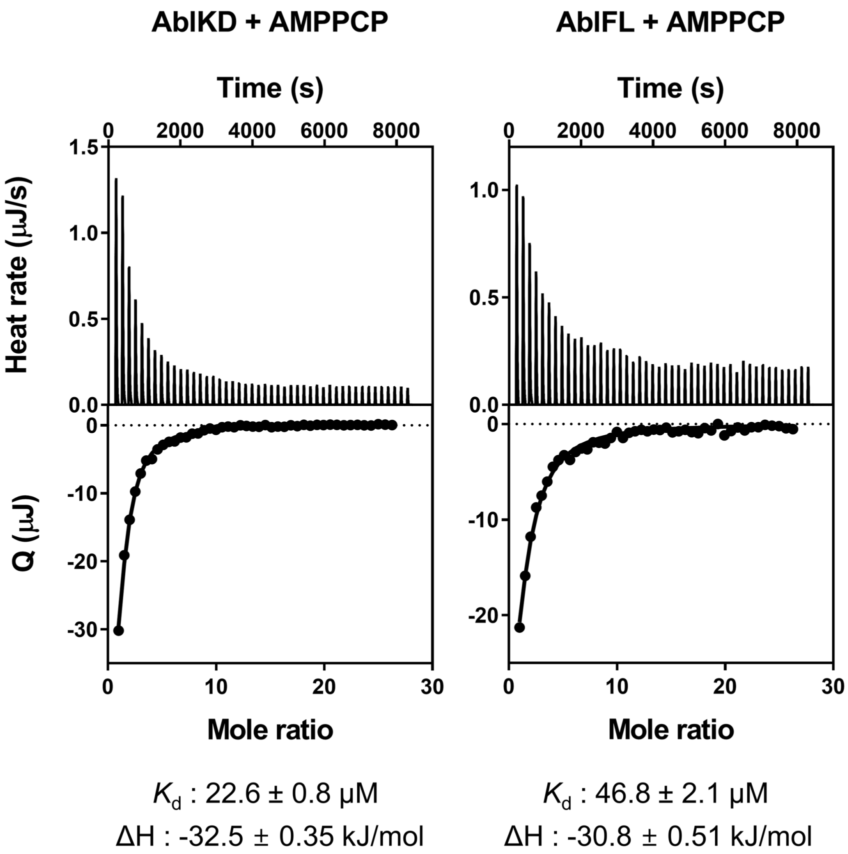


**Figure S11: ITC profiles for binding of AMPPCP to AblFL and AblKD.** Observed dissociation constants, $K_d$, and change in enthalpy, $\Delta H$, are shown with errors that represent 68.3% confidence interval of the fit of the data (± 1 s.d.). We find AMPPCP binds about 2-fold tighter to AblKD than AblFL, confirming that AMPPCP is shifting the open/closed equilibrium towards the open state.

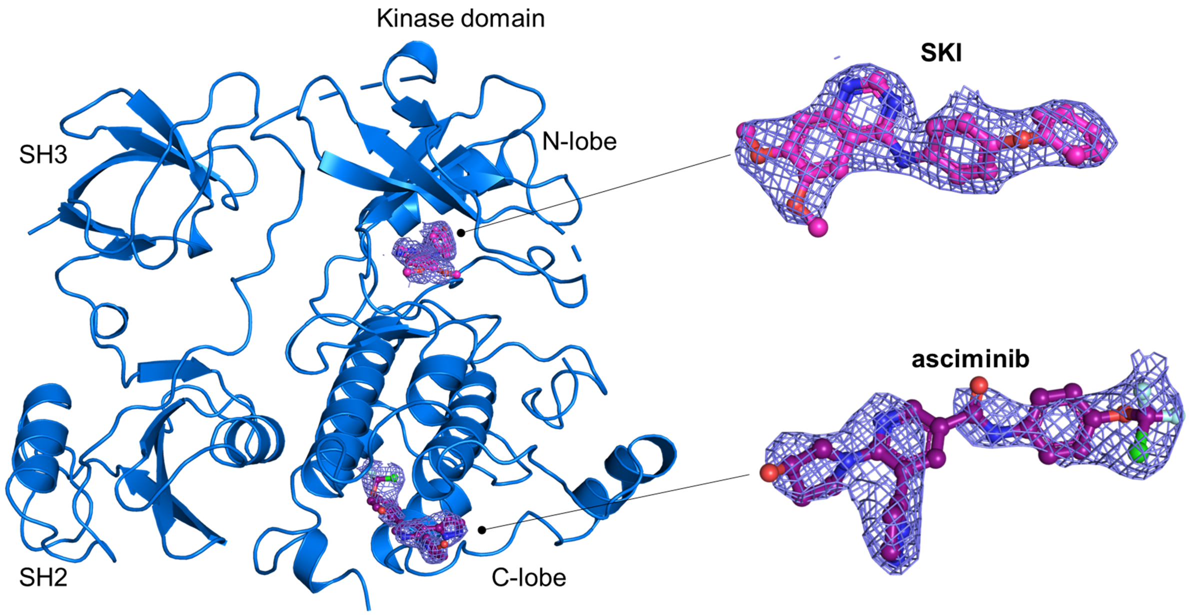


**Figure S12: 2m$F_o$-D$F_c$ electron density for SKI and asciminib in X-ray structure of AblFL-SKI-asciminib.** 2m$F_o$-D$F_c$ maps were contoured at 1σ (blue mesh). AblFL is represented as a cartoon (blue). SKI and asciminib are represented as spheres and sticks. Carbon, nitrogen, oxygen, chlorine and fluorine atoms are colored magenta, blue, red, green and mint, respectively.

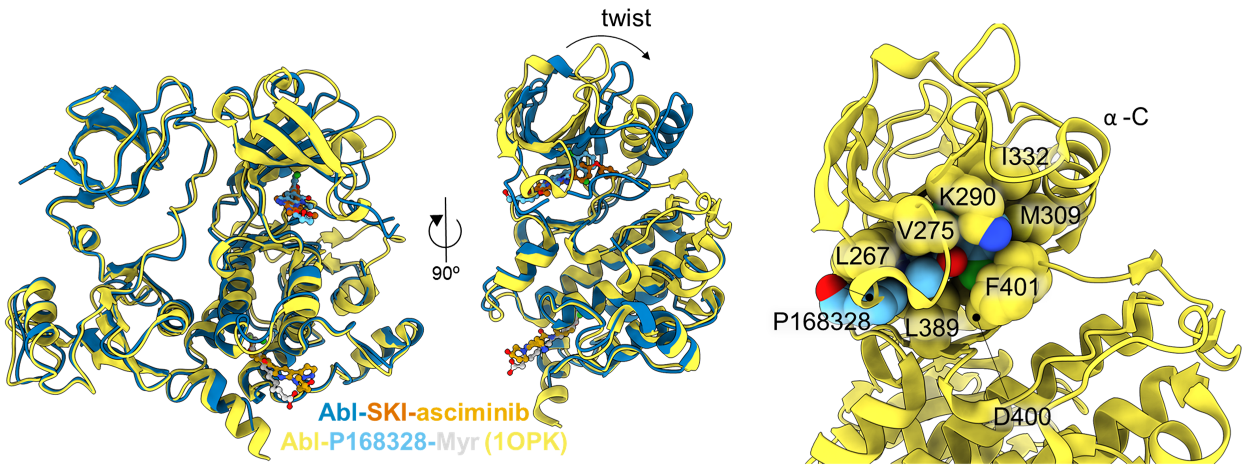


**Figure S13: Superposition of X-ray structures of Abl-SKI-asciminib (blue) and Abl-PD166326-myristate (yellow) (PDB-ID: 1OPK, PD166326 is an orthosteric inhibitor).** Due to the SKI-induced twist of the N-lobe, α-C helix in Abl-SKI-asciminib structure is shifted to "out" in comparison to α-C helix "in" for Abl-PD166326-myristate structure (4, 5). Structurally, we find PD166326 to be not as tightly packed as SKI (compare right panel with Fig. 5*C*). Oxygen, nitrogen and chlorine atoms are colored in red, blue and green, respectively. Carbon atoms are colored in dark orange, orange, light blue and grey for SKI, asciminib, P168328 and myristate, respectively.

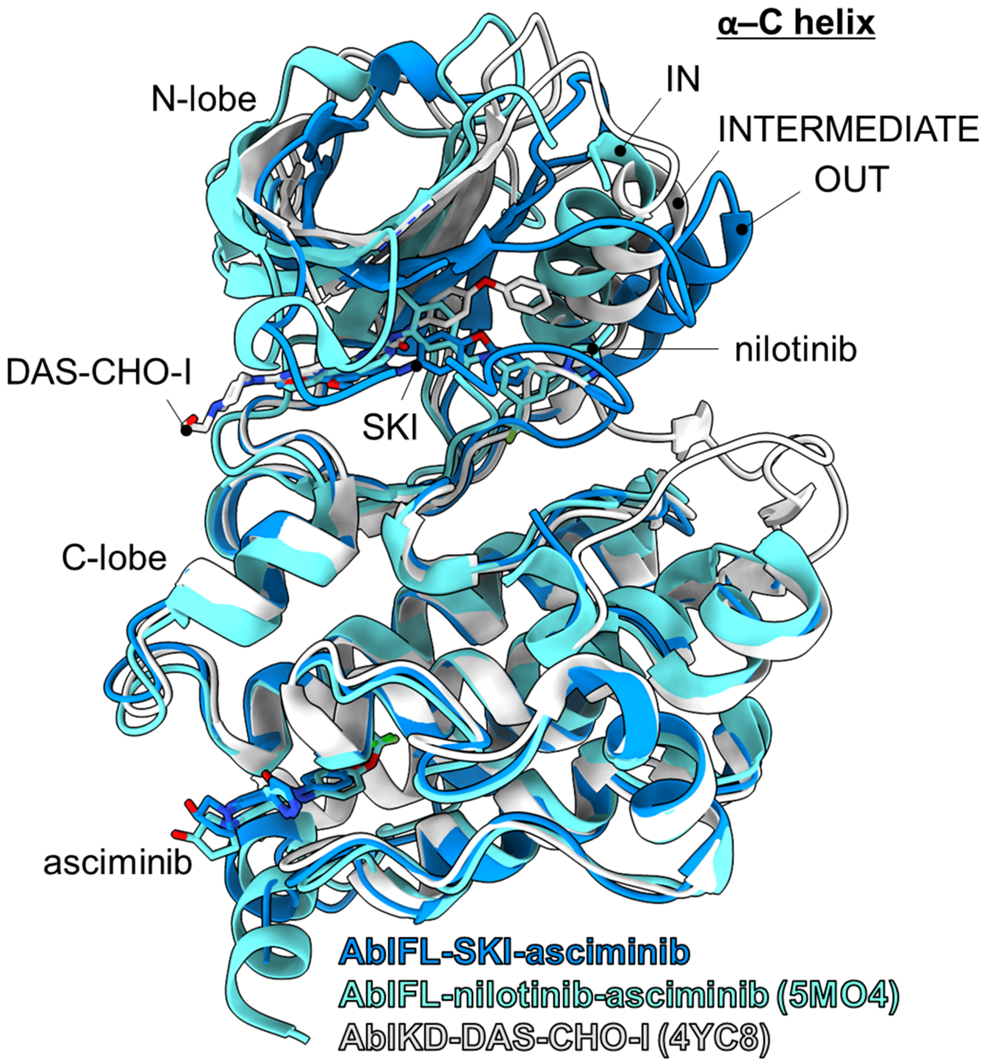


**Figure S14: Overlay of X-ray crystal structures for Abl-SKI-asciminib, Abl-nilotinib-asciminib (PDB-ID: 5MO4) and AblKD-DAS-CHO-I (PDB-ID: 4YC8) (5, 6).** Only catalytic domains are shown for clarity here. While Soellner and colleagues claimed that AblKD-DAS-CHO-I is the first α-C helix "out" inhibitor-bound Abl structure (6), we find that its α-C helix as intermediate state compared to α-C helix "in" in Abl-nilotinib-asciminib structure (5) and α-C helix "out" in our Abl-SKI-asciminib structure. AblKD-DAS-CHO-I was a structure of only the AblKD, not AblFL. Abl-SKI-asciminib is the first fully-closed structure of Abl, and SKI is an orthosteric inhibitor favoring the fully α-C helix "out", and closed conformation of Abl.

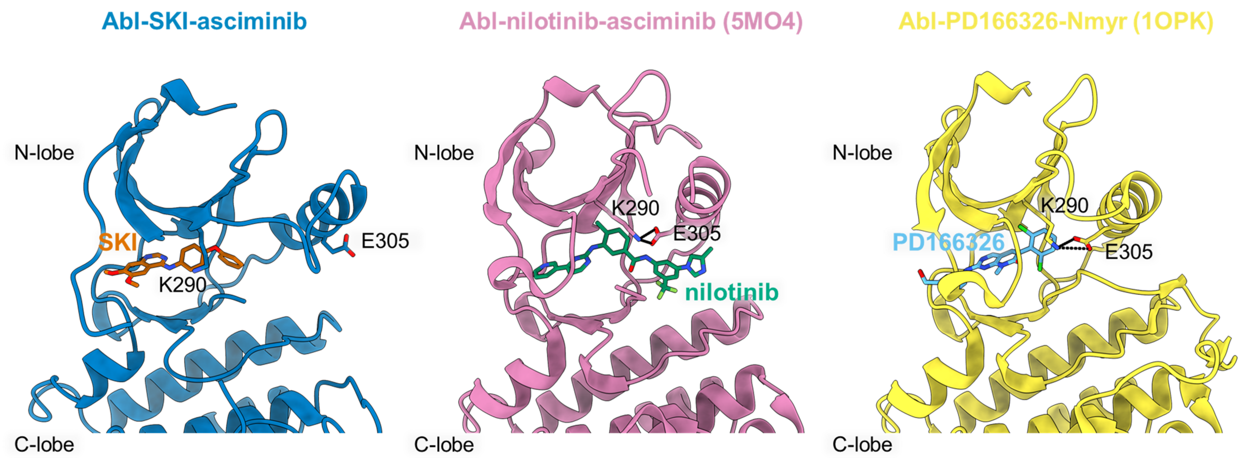


**Figure S15: Comparison of AblFL ternary X-ray structures highlighting canonical salt-bridge (K290-E305).** Only AblFL in complex with SKI and asciminib show broken canonical salt-bridge due to α-C helix in the "out" position, while the salt-bridge is established, with α-C helix in the "in" position, in the other closed ternary AblFL structures. This highlights the example of a truly fully-closed AblFL structure when in complex with SKI and asciminib.

**Table 1: X-ray structures data collection and refinement statistics**

Statistics for the highest-resolution shell are shown in parentheses.

| | AurA_danusertib_Mb1 | AurA_danusertib_Mb2 | Abl_SKI_asciminib |
|---|---|---|---|
| **Resolution range** | 45.4 - 2.6 (2.693 - 2.6) | 44.19 - 1.974 (2.044 - 1.974) | 46.56 - 2.865 (2.967 - 2.865) |
| **Space group** | I 2 2 2 | P 21 21 21 | P 21 21 21 |
| **Unit cell** | 75.517 90.805 143.13 90 90 90 | 65.619 73.225 179.315 90 90 90 | 92.478 103.213 107.963 90 90 90 |
| **Total reflections** | 110014 (11115) | 557749 (52621) | 48659 (4735) |
| **Unique reflections** | 15516 (1519) | 59706 (6010) | 24343 (2372) |
| **Multiplicity** | 7.1 (7.3) | 9.3 (8.8) | 2.0 (2.0) |
| **Completeness (%)** | 99.75 (99.35) | 95.64 (99.24) | 98.28 (89.70) |
| **Mean I/sigma I** | 8.74 (1.97) | 20.24 (2.07) | 4.93 (0.51) |
| **R-merge** | 0.1933 (1.341) | 0.0803 (1.118) | 0.08614 (1.367) |
| **CC1/2** | 0.993 (0.871) | 0.999 (0.731) | 0.997 (0.31) |
| **CC*** | 0.998 (0.965) | 1 (0.919) | 0.999 (0.688) |
| **Reflections used in refinement** | 15495 (1519) | 58998 (6010) | 24343 (2133) |
| **Reflections used for R-free** | 1068 (110) | 2924 (299) | 1969 (175) |
| **R-work** | 0.2057 (0.3628) | 0.1897 (0.3676) | 0.2922 (0.4674) |
| **R-free** | 0.2722 (0.4009) | 0.2303 (0.3483) | 0.3461 (0.4809) |
| **CC(work)** | 0.924 (0.735) | 0.925 (0.669) | 0.935 (0.415) |
| **CC(free)** | 0.912 (0.717) | 0.924 (0.696) | 0.929 (0.463) |
| **Number of non-hydrogen atoms**<br>Macromolecules<br>Ligands<br>Solvent | **3055**<br>2874<br>69<br>112 | **6140**<br>5587<br>85<br>468 | **6860**<br>6664<br>146<br>60 |
| **Protein residues** | 352 | 689 | 834 |
| **RMS(bonds)** | 0.008 | 0.011 | 0.006 |
| **RMS(angles)** | 1.23 | 1.44 | 1.05 |

| | | | |
|---|---|---|---|
| **Ramachandran favored (%)** | 96.55 | 97.93 | 98.52 |
| **Ramachandran allowed (%)** | 2.87 | 1.92 | 1.48 |
| **Ramachandran outliers (%)** | 0.57 | 0.15 | 0.00 |
| **Rotamer outliers (%)** | 2.58 | 2.14 | 0.41 |
| **Clashscore** | 4.29 | 4.76 | 44.65 |
| **Average B-factor** | **29.86** | **25.97** | **85.36** |
| Macromolecules | 28.49 | 23.75 | 85.56 |
| Ligands | 64.98 | 50.02 | 91.99 |
| Solvent | 43.24 | 48.18 | 38.33 |
